# Ikaros family proteins regulate developmental windows in the mouse retina through convergent and divergent transcriptional programs

**DOI:** 10.1101/2021.12.01.470829

**Authors:** Awais Javed, Pierre Mattar, Allie Cui, Michel Cayouette

## Abstract

Temporal identity factors regulate the competence of neural progenitors to generate specific cell types in a time-dependent manner, but how they operate remains poorly defined. In the developing mouse retina, the Ikaros zinc finger transcription factor Ikzf1 regulates the production of early-born cell types, except cone photoreceptors. In this study we show that Ikzf4, another Ikaros family protein, cooperates with Ikzf1 to control cone photoreceptor production during early stages of retinal development, whereas at late stages, when Ikzf1 is no longer expressed in progenitors, Ikzf4 is instead required for Müller glia production. Using CUT&RUN sequencing, we find that both Ikzf1 and Ikzf4 generally bind to the same genes involved in cone development and other early-born fates, but at different cis-regulatory elements. In late-stage progenitors, Ikzf4 re-localizes to bind target genes involved in Müller glia development and regulate their expression. Specifically, we show that Ikzf4 maintains Hes1 expression in differentiating cells using two Ikzf GGAA binding sites at the *Hes1* promoter, thereby favouring Müller glia fate commitment. These results uncover a combinatorial role for Ikaros family members in nervous system development and provide mechanistic insights on how they temporally regulate cell fate output.

## INTRODUCTION

Generation of cell diversity in the central nervous system (CNS) is a highly controlled and regulated process. Neural progenitors alter their potential to generate specific neurons and glia using both spatial and temporal patterning cues (Sagner and Briscoe, 2019). In the *Drosophila* nervous system, temporal patterning is regulated by the expression of transcription factors referred to as “temporal identity” factors, which control the developmental competence of neural progenitor cells, allowing cell-type production to change as development proceeds (Brody and Odenwald, 2000; Cleary and Doe, 2006; Erclik et al., 2017; Grosskortenhaus et al., 2005; Grosskortenhaus et al., 2006; Isshiki et al., 2001; Kambadur et al., 1998; Li et al., 2013; Novotny et al., 2002; Pearson and Doe, 2003). A classic example of temporal patterning in vertebrates is the developing mouse retina, where seven broad cell types are formed in sequential but overlapping manner from multipotent retinal progenitor cells (RPCs) (Rapaport et al., 2004; Young, 1985a, b). Retinal ganglion cells (RGCs), amacrine cells, cone photoreceptors and horizontal cells are mostly generated during the embryonic period of retinogenesis, whereas rod photoreceptors, bipolar cells, and Müller glia are primarily generated during the postnatal period (Carter-Dawson and LaVail, 1979a, b; Rapaport et al., 2004; Turner et al., 1990; Young, 1985a, b).

How exactly RPCs change competence over time to control retinal histogenesis remains poorly understood, although some progress was made in recent years (Davis et al., 2011; Decembrini et al., 2009; Dupacova et al., 2021; Georgi and Reh, 2010; Gordon et al., 2013; Iida et al., 2011; La Torre et al., 2013; Yang et al., 2003; Zibetti et al., 2019). Homologs of *Drosophila* temporal identity factors were found to regulate temporal patterning in mouse RPCs (Elliott et al., 2008; Javed et al., 2020; Mattar et al., 2015). Specifically, it was found that *Ikzf1* (*hunchback*), a member of the Ikaros family of zinc finger transcription factors, is necessary and sufficient to confer competence to generate most early-born cell types, except cone photoreceptors (Elliott et al., 2008). More recently, the homolog of *Drosophila pdm/nub*, *Pou2f1*, was shown to play a part in the temporal regulation of cone photoreceptor production by upregulating Pou2f2, which in turn represses the rod determinant Nrl in photoreceptor precursors, thereby favouring the cone fate (Javed et al., 2020). Intriguingly, Ikzf1 was found to upregulate Pou2f1 expression, but because Ikzf1 knockout retinas have normal numbers of cones, these results suggest that other, still unidentified factor(s), cooperate with Ikzf1 in RPCs to confer competence to generate cones (Elliott et al., 2008).

Mechanisms regulating the production of late-born cell types also remain incompletely understood. The homolog of *Drosophila castor*, *Casz1*, another zinc finger transcription factor, confers competence to generate late-born rod and bipolar cells in the mouse retina, but suppresses the production of Müller glia - the latest-born cell type (Mattar et al., 2015). Similarly, while Foxn4 was found to operate downstream of Ikzf1 and upstream of Casz1 to regulate mid-late temporal identity (Liu et al., 2020), it is not involved in Müller glia production. Instead, the temporal regulation of Müller glia production in the developing mouse retina is largely achieved by Nfi family of transcription factors (Clark et al., 2019), which are required to promote glial differentiation genes, even when expressed in early-stage RPCs that do not normally generate glia (Lyu et al., 2021). It remains unknown, however, how exactly Nfi factors are upregulated during late stages of retinogenesis to control glia production. Additionally, Sox9, Vsx2 and Sox2 are sufficient and required for Müller glia fate determination, but as they are expressed throughout retinal development, it is unclear how their pro-glial activities are gated (Lin et al., 2009; Livne-bar et al., 2006; Poche et al., 2008; Surzenko et al., 2013; Taranova et al., 2006).

There are five members of the Ikaros family of transcription factors in mice, which are homologous to the *Drosophila hunchback* gene, four of which are expressed in the developing retina (Elliott et al., 2008). Aside from Ikzf1, however, the role of Ikaros family members in the mammalian CNS has not been explored. We therefore wondered whether other Ikaros family members might contribute to regulate RPC competence during mouse retinogenesis.

We show here that Ikzf4 cooperates with Ikzf1 during early stages of retinogenesis to regulate cone development. Using Cleavage Under Target & Release Under Nuclease (CUT&RUN) sequencing, we report that Ikzf1 and Ikzf4 bind to different regulatory elements associated with the same sets of early cell type differentiation genes, including cone genes. While Ikzf1 expression is turned off at later stages of retinal development, Ikzf4 expression is maintained and regulates Müller glia development. Mechanistically, we find that Ikzf4 binds and regulates expression of several gliogenic genes, such as Notch targets and Nfia/Nfib. More specifically, we show that Ikzf4 binds to two specific ‘GGAA’ Ikaros binding motifs in the *Hes1* promoter that are essential for its regulatory activity. These results provide evidence of combinatorial roles for Ikzf1 and Ikzf4 during retinogenesis that regulates cell fate in a context- and temporal-dependent manner.

## RESULTS

### Ikzf4 is expressed in retinal progenitor cells at all stages of retinogenesis

We previously reported expression of Ikzf4 transcripts in the retina using in situ hybridization (Elliott et al., 2008), but the protein expression pattern remained unknown. To fill this knowledge gap, we characterized a commercially-available anti-Ikzf4 antibody. First, we electroporated CAG:GFP or CAG:Ikzf4-IRES-GFP vectors in E17 mouse retinas and immunostained retinal sections two days later with the Ikzf4 antibody. As expected, we found that the antibody recognizes overexpressed Ikzf4 proteins (Fig. S1A-C’). Second, to ensure that the antibody is specific for Ikzf4, we immunostained retinas isolated from *Ikzf4^+/+^* and *Ikzf4^-/-^* (RIKEN Bioresource <u>RRID: IMSR_RBRC06808)</u> mouse embryos at E12, E15 and P9. We found that Ikzf4 is expressed in virtually all retinal cells at E12 and E15, whereas it is restricted to a few retinal cell subtypes at P9 (Fig. S1D-H). No immunostaining signal was detected in the *Ikzf4^-/-^* retinas, indicating that the antibody specifically recognizes Ikzf4 (Fig. S1D-H’). Based on this expression pattern, we suspected that Ikzf4 was expressed in RPCs. Accordingly, we found that virtually all proliferating Ki67^+ve^ RPCs also stained for Ikzf4 at embryonic stages (Fig. 1A-D), and many Ki67^+ve^ cells also co-labelled with Ikzf4 at P0 and P2 (Fig. 1E-F), indicating that Ikzf4 is expressed in RPCs throughout development.

**Figure 1:**
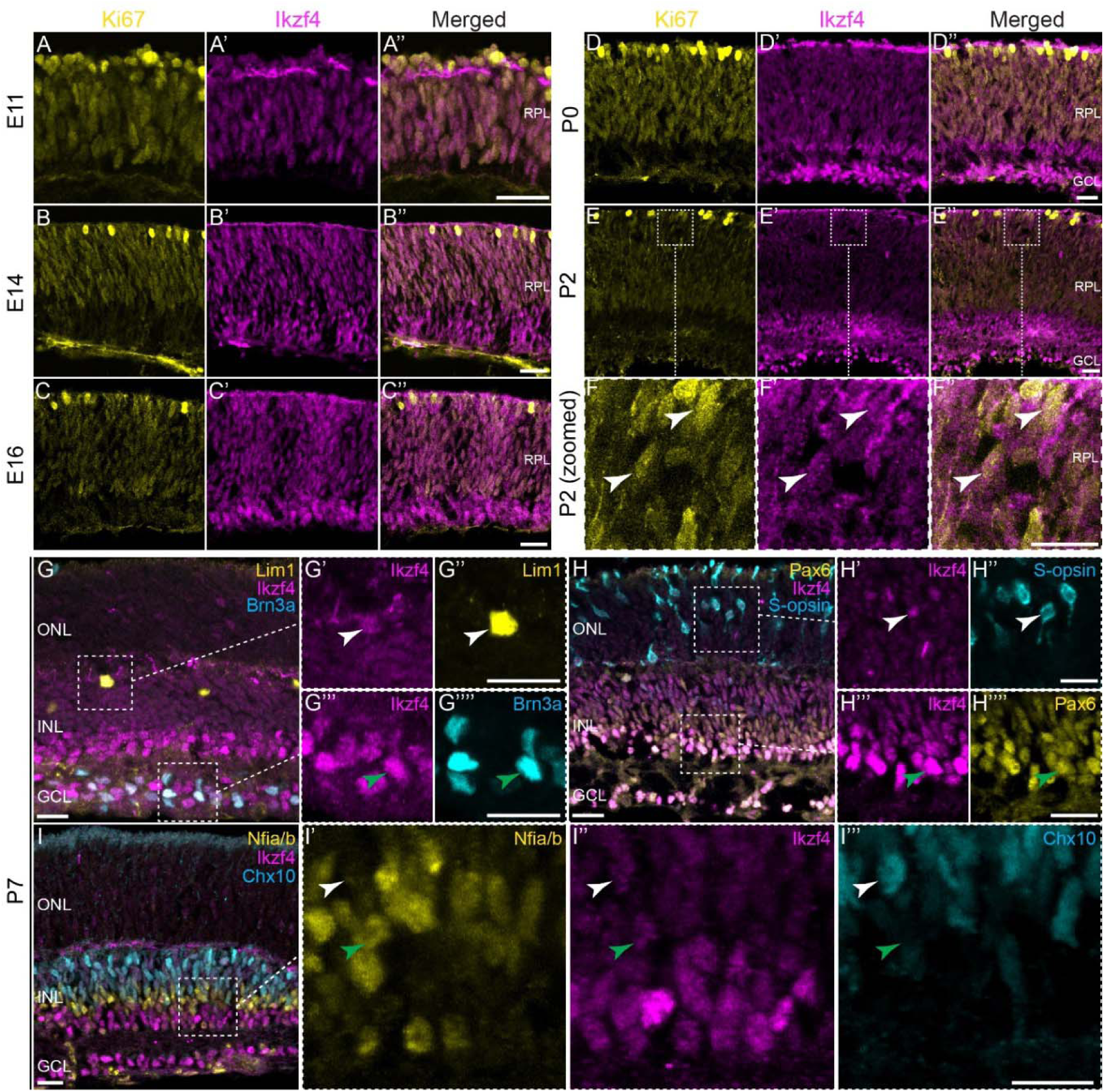
Ikzf4 is expressed during early and late retinogenesis. (A-E”) Co-immunostaining for Ki67 (yellow) and Ikzf4 (magenta) at various stages of mouse retinal development. (F-F”) Zoomed-in images of (E-E”), arrows show co-expression of Ki67 (yellow) and Ikzf4 (magenta) in some cells. (G-I’’’) Zoomed-out (G-I) and -in (G’-G’’’’,H’-H’’’’, I’-I’’’) examples of P7 mouse retinas co-immunostained for Ikzf4 (G’,G’’’,H’,H’’’,I’’), Lim1 (G’’), Brn3a (G’’’’), S-opsin (H’’), Pax6 (H’’’’), Nfia/b (I’) and Chx10 (I’’’). White arrows indicate Ikzf4^+ve^Lim1^+ve^ (G’-G’’), Ikzf4^+ve^S-opsin^+ve^ (H’-H’’) and Nfia/b^-ve^Ikzf4^+ve^Chx10^+ve^ cells. Green arrows indicate Ikzf4^+ve^Brn3a^+ve^ (G’’’-G’’’’), Pax6^+ve^Ikzf4^+ve^(H’’’-H’’’’) and Nfia/b^+ve^Ikzf4^+ve^Chx10^-ve^ (I’-I’’’) Zoomed in regions are highlighted with dashed boxes. RPL: Retinal progenitor layer. ONL: Outer nuclear layer. INL: Inner nuclear layer. GCL: Ganglion cell layer. Scale bars: 20μm (A-E’’), 10μm (F-I’’’).

We next compared our immunostaining data for Ikzf4 with published scRNA-seq datasets in the developing mouse and human fetal retinas (Clark et al., 2019; Lu et al., 2020). Focusing on the RPC population, we generated UMAP plots by sub-setting the RPC populations at various developmental stages. We found that *Ikzf4*/*IKZF4* mRNA is expressed in both early and late RPCs (Fig. S1I-J), consistent with our immunostaining results. In addition to RPCs, we found *Ikzf4/IKZF4* expression in some differentiated retinal cell type clusters like rods, amacrine, horizontal, and retinal ganglion cells (Fig. S2A-B). This suggests that Ikzf4 is expressed in the mouse and human retina throughout development.

To assess expression of Ikzf4 in mature cells, we used cell-type specific antibodies at P7. We detected expression of Ikzf4 in early-born cell types such as Lim1^+ve^ horizontal cells, Brn3a^+ve^ RGCs, Pax6^+ve^ amacrine cells and S-opsin^+ve^ cone photoreceptors (Fig. 1G-H’’’’). We could also detect weak Ikzf4 immunostaining in S-opsin^-ve^ cells in the ONL, suggesting some expression in rod photoreceptors (Fig.1 H’-H’’). These results are consistent with our *Ikzf4*/*IKZF4* expression analysis in scRNA-seq datasets (Clark et al., 2019; Lu et al., 2020) (Fig. S2A-B). Interestingly, we also found Ikzf4 expression in Nfia/b^+ve^ cells that are either Chx10^-ve^ or Chx10^+ve^ (Fig. 1I-I’’’), likely representing Müller glia and bipolar cells, respectively. Taken together, these data indicate that Ikzf4 is detected in all early-born cell types and in some late-born cell types.

### Ikzf4 overexpression in late-stage retinal progenitors promotes immature cone and Müller glia production

We next investigated the function of Ikzf4 in the developing mouse retina. We first misexpressed Ikzf4 in late-stage RPCs and asked if this would be sufficient to induce early-born cell type production. We infected P0 retinal explants with retroviral vectors expressing Venus or Ikzf4-IRES-Venus, and analyzed individual clone compositions 14 days later. We used Rxrg staining to identify cone photoreceptors, and cell morphology combined with nuclear layer position to identify other cell types, as we did previously (Elliott et al., 2008; Javed et al., 2020). We found many Venus^+ve^/Rxrg^+ve^ cells located in the ONL in Ikzf4-IRES-Venus infected clones (Fig. 2A-C’), suggesting that Ikzf4 is sufficient to promote cone photoreceptor production outside their normal window of development. Consistently, we observed an increase in the proportion of Venus^+ve^ cells that co-stained for Rxrg in Ikzf4-infected retinas, as well as an increase in Müller glia (Fig. 2D, E), which was accompanied by a decrease in rods, both as a proportion of all cells analyzed and as number of rods per clone (Fig. 2F). Reinforcing this finding, we observed a reduction in *Nrl* and *Nr2e3* transcript levels, two rod-specific genes, and an increase in *Rxrg* expression in the GFP^+ve^ cell population 6 days after electroporation of CAG:Ikzf4-IRES-GFP in P0 retinal explants (Fig. 2G). The number of GFP^+ve^ cells in the ONL that stain for Nrl and Nr2e3 was also reduced following Ikzf4 expression (Fig. 2H-N).

**Figure 2:**
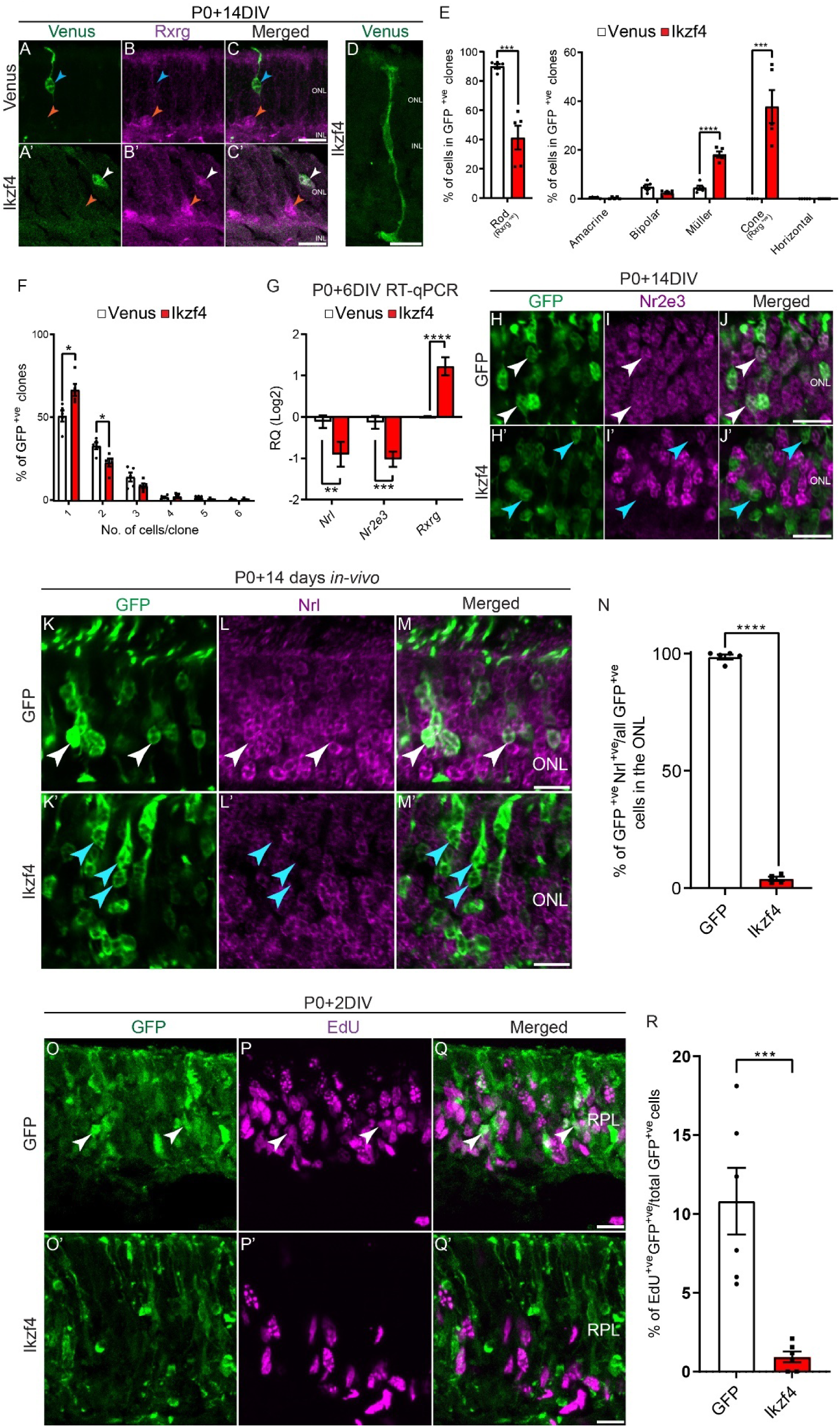
Ikzf4 promotes cones and Müller glia fate specification from late-stage RPCs. (A-C) Examples of clones obtained after infection with retroviral vectors expressing Venus (A-C) and Ikzf4-IRES-Venus (A’ -C’ ) co-immunostained with Rxrg (B-B’ ), a cone marker. (D) Example of Müller glia generated by Ikzf4 retroviral infection. (E-F) Retroviral lineage analysis of Venus control (469 clones counted) and Ikzf4-IRES-Venus (388 clones counted) overexpression in late-stage retinas. (E) Quantifications for cell type analysis was based on morphology and laminar positioning of the cell bodies in the retina (Venus, n=5; Ikzf4, n=5). Cones (Rxrg^+ve^) and rods (Rxrg^-ve^) were counted in the ONL based on Rxrg expression. (F) Quantifications of the number of cells per clone for the data presented in (E). (G) RT-qPCR analysis of *Nrl*, *Nr2e3* and *Rxrg* expression from sorted GFP^+ve^ cells 6 days after electroporation of P0 retinal explants with either GFP (n=5) or Ikzf4 (n=5). (H-J’) Examples of retinal explants electroporated at P0 with either GFP (H-J) or Ikzf4-IRES-GFP (H’ -J’ ) and immunostained for Nr2e3 (I-I’ ) 14 days later. White arrows represent electroporated cells positive for Nr2e3 and cyan arrow represent electroporated cells without Nr2e3 immunostaining. (K-M’ ) Examples of retinas electroporated *in vivo* with either GFP (K-M) or Ikzf4 (K’ -M’ ) and immunostained for Nrl (L-L’ ) 14 days after electroporation. White arrows label GFP^+ve^Nrl^+ve^ cells; cyan arrows label GFP^+ve^Nrl^-ve^ cells. (N) Quantification of the number of GFP^+ve^Nrl^+ve^ cells (GFP, n=5; Ikzf4, n=4). (O-Q’) Examples of P0 retinal explants electroporated with either GFP (O-Q) or Ikzf4 (O’ -Q’ ), with 30μM EdU added to the culture medium 2 days later. Retinal explants were immunostained with EdU after fixation. White arrows indicate GFP^+ve^EdU^+ve^ cells. (R) Quantifications of the number of EdU^+ve^GFP^+ve^ cells (GFP (593 cells counted), n=6; Ikzf4 (676 cells counted), n=6). *p<0.05, **p<0.01, ***p<0.001, ****p<0.0001. Statistics: Two-tailed unpaired t-test (E-F, N, R), Mann-Whitney test (G). RPL: Retinal progenitor layer. ONL: Outer nuclear layer. INL: Inner nuclear layer. RQ: Relative quantitation. Scale bars: 10μm (A-D, H-J’, K-M’, O-Q’).

As we found that the number of small clones is increased following Ikzf4 expression (Fig. 2F), we postulated that Ikzf4 might promote precocious cell cycle exit or cell death. To distinguish between these possibilities, we electroporated CAG:GFP or CAG:Ikzf4-IRES-GFP vectors in P0 retinas. After culturing retinal explants for 2 days, we added EdU for 2 hours then fixed and stained the explants for GFP and EdU. We observed a significant decrease in EdU^+ve^GFP^+ve^ cells after expression of Ikzf4 (Fig. 2O-R). As we found no change in the number of cells staining for cleaved Caspase-3 (Fig. S3A-C’), we conclude that Ikzf4 expression reduces clone size by promoting early cell cycle exit, rather than cell death. Finally, we also found that most 1- and 2-cell clones in the Ikzf4 condition contain either cones or Müller glia, whereas control clones contain mostly rods (Fig. S3D-E). Together, these results suggest that Ikzf4 is sufficient to promote cones and Müller glia production at the expense of rods when expressed in late RPCs.

To buttress the retroviral lineage tracing results and provide more in-depth analysis of the cell types produced, we electroporated retinal explants with either CAG:GFP or CAG:Ikzf4-IRES-GFP at P0 and determined the identity of GFP^+ve^ cells 14 days later using immunostaining for cell-type specific markers. As observed with retroviral vectors, we found that Ikzf4- electroporated retinas contain more Rxrg^+ve^ cells in the ONL compared to the control CAG:GFP (Fig. S3F-I). Intriguingly, however, the GFP^+ve^Rxrg^+ve^ cells did not stain for the mature cone markers S-opsin, M-opsin or PNA (Fig. S3J-M’). We therefore wondered whether these GFP^+ve^Rxrg^+ve^ cells might be mis-localized retinal ganglion cells (RGCs), which also express Rxrg (Mori et al., 2001). We co-immunostained sections of Ikzf4-electroporated retinas with GFP, Rxrg, and the RGC markers Brn3a and Brn3b. We found that Ikzf4 induces production of Rxrg^+ve^ cells that do not express Brn3a or Brn3b, indicating they are not RGCs (Fig. S3N-Q’). Instead, Ikzf4-expressing GFP^+ve^ cells in the ONL co-labelled with Crx/Otx2, which specifically labels photoreceptor cells in this layer (Fig. S3R-T’). Thus, overexpression of Ikzf4 in late-stage RPCs is sufficient to induce production of cone photoreceptors that remain incompletely differentiated.

Similar to what we observed with retroviral vectors, we observed an increase in Müller glia production after overexpression of Ikzf4-IRES-GFP at P0, as determined by the GFP^+ve^Sox2^+ve^ cells, GFP^+ve^Hes1^+ve^Chx10^-ve^ and GFP^+ve^Lhx2^+ve^Nfia/b^+ve^ cells found in the INL (Fig. S3U-Y’). These results indicate that, in addition to immature cones, Ikzf4 promotes production of Müller glia when expressed in late-stage RPCs.

### Ikzf4 is required for Müller glia development

Based on the above results, we hypothesized that Ikzf4 may be required for cone and Müller glia development. We therefore analyzed retinas from *Ikzf4^+/+^*, *Ikzf4^+/-^*, and *Ikzf4^-/-^* mice and quantified the various retinal cell types using specific markers at P10, a stage when cell genesis is largely complete. While we found no change in the number of RGCs (Brn3a^+ve^), amacrine cells (Pax6^+ve^), horizontal cells (Lim1^+ve^) or bipolar cells (Otx2^+ve^), we observed a reduction in Sox2^+ve^ and Lhx2^+ve^ cells in the INL of *Ikzf4^-/-^* retinas compared to *Ikzf4^+/+^* and *Ikzf4^+/-^* retinas (Fig. 3A-C). Surprisingly, we found no difference in the number of Rxrg^+ve^ cells in the ONL in *Ikzf4^-/-^* retinas compared to control *Ikzf4^+/+^* (Fig. 3C). These results indicate that Ikzf4 is required for Müller glia development, but dispensable for cone photoreceptor production, even though it is sufficient to induce immature cone production from late RPCs.

**Figure 3:**
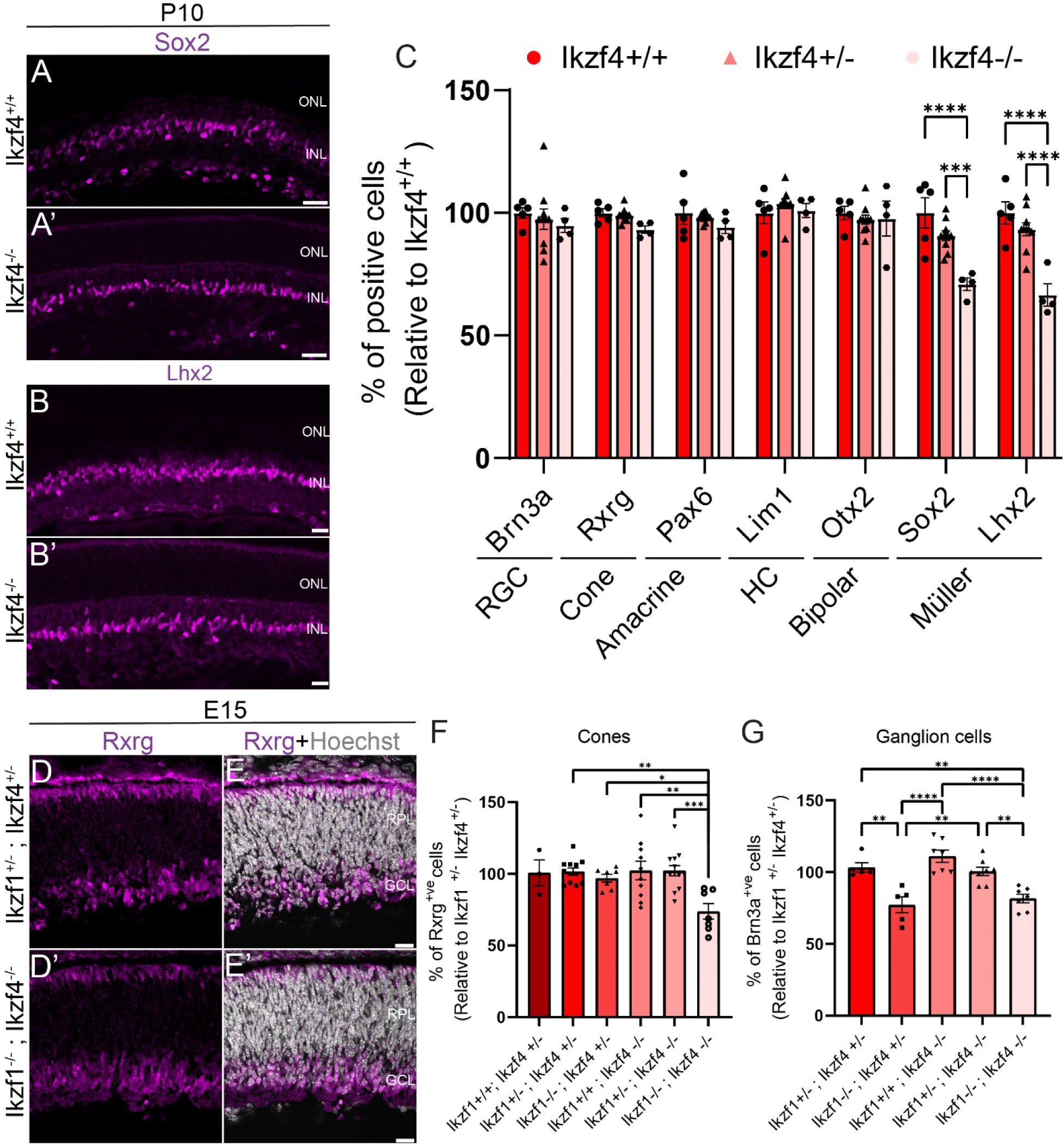
Combinatorial requirement for Ikzf1 and Ikzf4 in the production of cones during early retinogenesis and Müller glia during late retinogenesis. (A-C) Examples of Sox2 (A-A’) and Lhx2 (B-B’) immunostaining in either Ikzf4^+/+^ (A-B) or Ikzf4^-/-^ (A’-B’) mouse retinas at P10. (C) Quantifications of various retinal cell types using specific markers in either Ikzf4^+/+^ (n=7), Ikzf4^+/-^ (n=13) or Ikzf4^-/-^ (n=7) mouse retinas. (D-E’) Examples of Rxrg (D-D’) immunostaining in either Ikzf1^+/-^Ikzf4^+/-^(D-E) or Ikzf1^-/-^Ikzf4^-/-^(D’-E’) mouse retinas at E15. (F) Quantifications of Rxrg^+ve^ cells in either Ikzf1^+/+^Ikzf4^+/-^(n=3), Ikzf1^+/-^Ikzf4^+/-^(n=12), Ikzf1^-/-^ Ikzf4^+/-^(n=7), Ikzf1^+/+^Ikzf4^-/-^(n=10), Ikzf1^+/-^Ikzf4^+/-^(n=13) or Ikzf1^-/-^Ikzf4^-/-^(n=7) mouse retinas. (G) Quantifications of Brn3a^+ve^ cells in either Ikzf1^+/-^Ikzf4^+/-^(n=5), Ikzf1^-/-^Ikzf4^+/-^(n=5), Ikzf1^+/+^Ikzf4^-/-^ (n=7), Ikzf1^+/-^Ikzf4^+/-^(n=8) or Ikzf1^-/-^Ikzf4^-/-^(n=7) mouse retinas. *p<0.05, **p<0.01, ***p<0.001, ****p<0.0001. Statistics: One-way ANOVA with Tukey correction (C, F-G). RPL: Retinal progenitor layer. ONL: Outer nuclear layer. INL: Inner nuclear layer. GCL: Ganglion cell layer. Scale bars: 10μm (A-B’, D-E’).

### Ikzf4 cooperates with Ikzf1 to control cone photoreceptor development

We have previously shown that inactivation of Ikzf1 decreases early-born cell type production, with the exception of cone photoreceptors, which are unaltered in Ikzf1 KO mice (Elliott et al., 2008). As Ikzf1 and Ikzf4 are expressed in the same cells as early as E11 (Fig. S4A-D), we wondered whether Ikzf1 and Ikzf4 might genetically cooperate to regulate cone development. To test this idea, we generated double knockout mice of Ikzf1 and Ikzf4. Interestingly, when both alleles of *Ikzf1* were knocked out together with one or two alleles of *Ikzf4* (*Ikzf1^-/-^*;*Ikzf4^+/-^* and *Ikzf1^-/-^;Ikzf4^-/-^*), embryos were paler and exhibited reduced liver size compared to *Ikzf1^+/-^*;*Ikzf4^-/-^* and *Ikzf1^+/-^;Ikzf4^+/-^* embryos (Fig. S4E-H). Moreover, we found that *Ikzf1^-/-^;Ikzf4^+/-^* and *Ikzf1^-/-^;Ikzf4^-/-^* mice die at early postnatal stages, usually before P2, whereas single *Ikzf1^-/-^ or Ikzf4^-/-^* mice are viable, showing functional redundancy for survival. Therefore, we focused our analysis of the retina at embryonic stages.

We first quantified cone numbers using Rxrg immunostaining at E15, when cone genesis is at its peak. While we found no difference in cone photoreceptor numbers between most genotypes analyzed, we found a significant reduction in cone numbers in *Ikzf1^-/-^;Ikzf4^-/-^* double knockout animals (Fig. 3D-F), indicating that Ikzf1 and Ikzf4 cooperate to control cone photoreceptor development. We also asked whether Ikzf4 might function with Ikzf1 to regulate production of other early-born cell types, which are only mildly decreased in *Ikzf1* KO retinas (Elliott et al., 2008). As expression of specific amacrine and horizontal cell markers generally come on at postnatal stages (Clark et al., 2019) when double knockout animals are lethal, we focused our attention on RGC production, as Brn3b is robustly expressed at E15.5 (Xiang, 1998). We found that the number of RGCs was reduced in *Ikzf1^-/-^; Ikzf4^+/-^*, consistent with our previously-published data (Elliott et al., 2008), but this RGC loss was not exacerbated in *Ikzf1^-/-^ ;Ikzf4^-/-^* double knockout animals. Taken together, these results show that Ikzf1 and Ikzf4 cooperate to regulate cone, but not RGC development.

### Ikzf1 and Ikzf4 regulate genes involved in early-born cell type production

To explore how Ikzf1 temporally reprograms late-stage RPCs, we misexpressed Ikzf1 in P0 retinas and carried out CUT&RUN and RNA-sequencing, as schematized (Fig. 4A). In parallel, we performed CUT&RUN on E14 and P0 retinal extracts using a previously-validated anti-Ikzf4 antibody (S1A-H’). Using MACS2 peak calling on the CUT&RUN datasets, we found 1787 Ikzf1 peaks, 2472 Ikzf4 peaks at E14, and 2130 Ikzf4 peaks at P0 (Fig. S5A). When we intersected the peaks between the Ikzf1 and Ikzf4 E14 and P0 datasets using bedtools (Quinlan and Hall, 2010), we only found a few overlapping (Fig. S5A), suggesting that Ikzf1 and Ikzf4 do not bind to the same cis-regulatory elements (CREs) in RPCs. We next assessed the CUT&RUN feature distribution using ChIPSeeker to annotate the peaks (Yu et al., 2015). We found that Ikzf1 binding was mostly observed in intergenic and intronic regions, away from the promoters (Fig. S5B), similar to what was previously reported in B-cells (Schwickert et al., 2014). On the other hand, Ikzf4 binding, both at E14 and P0, was primarily observed at promoters. We further analyzed Ikzf1 and Ikzf4 binding peaks by performing Hypergeometric Optimization of Motif EnRichment (HOMER) to discover transcription factor motifs present in the dataset (Heinz et al., 2010). We found that the canonical Ikzf binding motif ‘GGAA’ or the complementary sequence is present in Ikzf1 and Ikzf4 CUT&RUN datasets (Fig. S5C), as previously reported for Ikzf family members (Molnár and Georgopoulos, 1994). Taken together, these data suggest that Ikzf1 binds to GGAA motifs in distal and intronic CRE, whereas Ikzf4 binds to GGAA motifs close to promoters.

**Figure 4:**
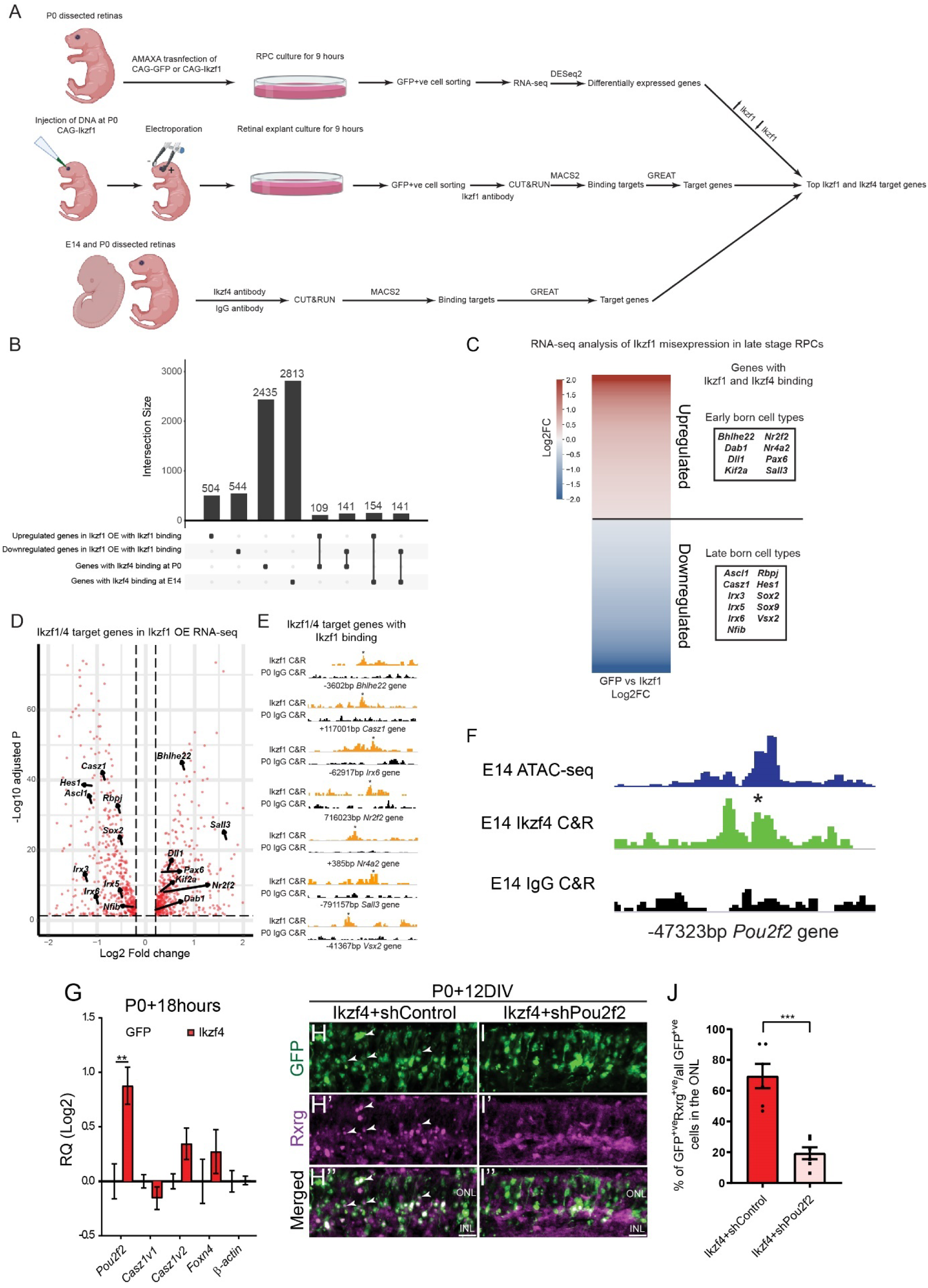
Ikzf1 and Ikzf4 bind to target genes involved in early-born cell type production. (A) Schematic showing how the experiment in (B-F) was performed. (B) Upset plot representing the overlap between up/downregulated genes following Ikzf1 OE with Ikzf1 binding and genes bound by Ikzf4 at E14 and P0. (C) Heatmap showing the top upregulated and downregulated genes in the RNA-seq analysis of Ikzf1 overexpression bound by Ikzf1 and Ikzf4 at either E14 or P0. (D) Volcano plot showing highlighted genes from (C) projected on the Ikzf1 RNA-seq data. (E) Genomic tracks of Ikzf1 CUT&RUN (orange) and P0 control IgG (black) on early and late genes found in (C-D). Asterisks indicate called peak. (F) Genomic tracks of ATAC-seq at E14 in blue (Aldiri et al. 2017), Ikzf4 CUT&RUN peaks at E14 in green, IgG Control CUT&RUN at E14 in black at genomic location of 47323bp upstream from the *Pou2f2* promoter. Asterisk indicates called peak. (G) RT-qPCR analysis *Pou2f2*, *Casz1v1*, *Casz1v2*, *Foxn4* or β *ctin* from sorted GFP^+ve^ cells 18hours after electroporation of P0 retinal explants with either GFP (n=5) or Ikzf4 (n=5). (H-I’’) Examples of retinal explants electroporated at P0 with Ikzf4-IRES-GFP and either shControl (H-H’’) or shPou2f2 (I-I’’) and immunostained for Rxrg (H’-I’). White arrows indicate GFP^+ve^Rxrg^+ve^ cells in the ONL. (J) Quantification of GFP^+ve^Rxrg^+ve^ cells in the ONL (Ikzf4+shControl, n=6; Ikzf4+shPou2f2, n=6). Statistics: Mann-Whitney test (E), Two-tailed unpaired t-test (H). RQ: Relative quantitation. ONL: Outer nuclear layer. INL: Inner nuclear layer. Scale bars: 10μm.

We next asked whether Ikzf1 and Ikzf4 might bind the same genes, but at different CREs. We first curated the top differentially expressed genes by comparing the RNA-seq datasets from the CAG-GFP and CAG-Ikzf1-transfected RPCs using DESeq2 (Fig. 4A, Supplementary file 1). We performed gene ontology (GO) enrichment using GONet to find the top biological processes affected by Ikzf1 misexpression (Pomaznoy et al., 2018). In the upregulated list, we found many genes associated with retinal ganglion cells and amacrine cell development, such as *Tfap2b*, *Pou4f1*, *Isl1*, *Dll1/4* and *Pax6* under the GO biological process “animal organ development”, as expected since Ikzf1 promotes these early-born cell fates (Fig. S5D, Supplementary file 2) (Elliott et al., 2008). Conversely, in the downregulated list, we found many genes associated with bipolar and Müller glia development such as *Hes1*, *Irx3*, *Notch1/3*, *Rbpj*, *Sox2*, *Sox8* and *Sox9* under the GO biological process “negative regulation of nervous system development” (Fig. S5D, Supplementary file 2). We also found the late temporal factor *Casz1* in the same GO biological process, consistent with our previously published results that Ikzf1 represses *Casz1* (Mattar et al., 2015). To narrow down the list of target genes, we next compared the list of significantly altered transcripts in Ikzf1 misexpression RNA-seq to that of bound regions in Ikzf1 CUT&RUN using Genomic Regions Enrichment of Annotations Tool (GREAT) (McLean et al., 2010). We found around 1000 total upregulated and downregulated transcripts that are bound by Ikzf1 (Fig. 4B). Out of these, retinal ganglion cell and amacrine cell development genes were enriched in the upregulated category, whereas bipolar cell and Müller glia development genes were enriched in the downregulated category, consistent with Ikzf1 functioning as an early temporal identity factor (Supplementary file 1).

We next compared Ikzf1 RNA-seq/CUT&RUN target gene list with the Ikzf4 CUT&RUN E14/P0 target gene list to find common genes regulated by both Ikzf factors. After performing peak to gene annotation using GREAT, we observed over 150 genes regulated and bound by Ikzf1 that are also bound by Ikzf4 (Fig. 4B). In the upregulated category, we observed genes associated with cone photoreceptor development also bound by Ikzf1 and Ikzf4, such as *Kif2a*, *Nr2f2*, and *Sall3* (de Melo et al., 2011; Kallman et al., 2020; Satoh et al., 2009), and amacrine/ganglion/horizontal cell development, such as *Bhlhe22*, *Nr4a2*, *Dll1*, *Pax6*, *Tfap2b* and *Dab1* (Feng et al., 2006; Jiang and Xiang, 2009; Jin et al., 2015; Marquardt et al., 2001; Rice and Curran, 2000; Riesenberg and Brown, 2016) (Fig. 4C). These results suggest that Ikzf1 and Ikzf4 have common target genes that control the production of cones and other early-born cell types, although they bind different CREs of these genes (Fig. S5A). Conversely, the downregulated category contained genes bound by both Ikzf1 and Ikzf4 and involved in late-born cell type production such as *Casz1*, *Irx3/5/6*, *Sox2/9*, *Nfib* and *Vsx2*, as represented by volcano plot for gene expression and IGV genomic tracks for CUT&RUN binding peaks (Fig. 4D-E). Together, these data suggest that Ikzf1/Ikzf4 both repress and promote expression of genes involved in late- or early-born cell production, respectively.

We previously showed that Ikzf1 upregulates *Pou2f1* when overexpressed in early-stage retina, but not when misexpressed in late-stage retinas (Javed et al., 2020). Pou2f1 in turn promotes *Pou2f2* expression, which then represses the rod-promoting transcription factor *Nrl* to favour cone development (Javed et al., 2020). Because Ikzf1 cooperates genetically with Ikzf4 to promote the cone fate, we wondered whether Ikzf4 might also upregulate *Pou2f1/2* expression. As we misexpressed Ikzf1 in late-stage retinas, we did not observe upregulation of *Pou2f1/2* in Ikzf1 RNA-seq or found peaks at *Pou2f1/2* in Ikzf1 CUT&RUN (data not shown). On the other hand, Ikzf4 CUT&RUN analysis at E14 revealed that Ikzf4 binds at a region of open chromatin 47kbp upstream of the *Pou2f2* promoter (but did not bind regulatory regions associated with *Pou2f1*), which was scored as a significant *Pou2f2*-associated peak by GREAT (Fig. 4F). To test whether Ikzf4 regulates *Pou2f2* expression, we electroporated P0 retinas with either CAG:GFP or CAG:Ikzf4-IRES-GFP and sorted GFP+ cells 18 hours later to perform RT-qPCR. We found that Ikzf4 upregulates transcript levels of *Pou2f2*, whereas there is no change in the expression of temporal identity factors such as *Casz1v1*, *Casz1v2,* and *Foxn4,* which are regulated by Ikzf1 (Fig. 4G) (Liu et al., 2020; Mattar et al., 2015). These results suggest that Ikzf4 promotes cone development, at least in part, by binding and upregulating *Pou2f2* gene expression, in addition to other genes discussed above. To test whether Ikzf4 requires *Pou2f2* to promote cone production, we co-electroporated Ikzf4-IRES-GFP with either shControl or shPou2f2 in P0 retinal explants and analyzed cones generated 12 days later. We found a significant decrease in the number of GFP^+ve^Rxrg^+ve^ cells in the ONL when Pou2f2 was knocked-down concomitantly with Ikzf4 expression (Fig. 4H-J). These results show that Ikzf4, similar to Pou2f1, requires *Pou2f2* to promote cone development, suggesting a transcriptional cascade in which Ikzf4 activates *Pou2f2* expression to repress *Nrl* and favour cone production.

### Ikzf4 binds and upregulates genes involved in Müller glia production

We next asked how Ikzf4 could regulate Müller glia production during late stages of retinogenesis. We first investigated stage-specific peaks for Ikzf4 at E14 and P0. We computed the overlapping binding peaks between E14 and P0 stages using bedtools. We labelled Ikzf4 binding peaks found only at the E14 stage as ‘E14 exclusive’, those found only at P0 stage as ‘P0 exclusive’, and those found at both E14 and P0 as ‘Common’ peaks. We noticed that, out of the total binding peaks at each stage, only 1010 peaks were ‘Common’ (Fig. 5A, Supplementary file 3). To identify the genes associated with the E14 exclusive, P0 exclusive and Common binding peaks, we performed GO classification using GREAT and annotated the top 5 GO biological processes (McLean et al., 2010). In the top 5 GO biological processes for all three categories, we found “Positive regulation of Notch signaling pathway” only in P0 exclusive peaks (Fig. 5A), suggesting a prominent role of Ikzf4 in the regulation of Notch signaling during late retinogenesis.

**Figure 5:**
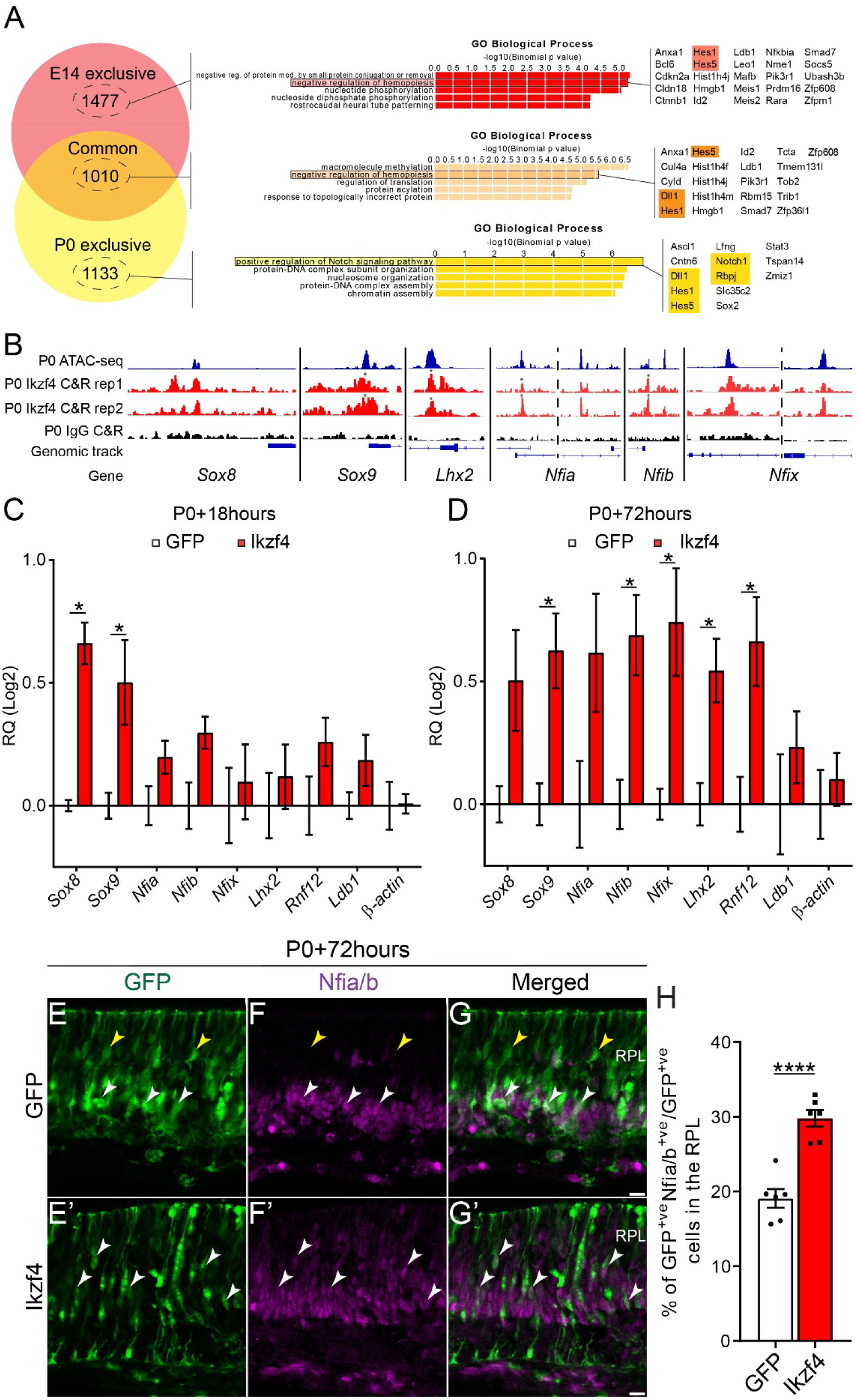
Ikzf4 binds and regulates expression of Müller specification genes. (A) Venn diagram representing the overlap of the number of Ikzf4 binding peaks between E14 and P0. Gene Ontology classification of the genes in proximity of Ikzf4 binding peaks. GO terms and associated notch signaling genes in the list are highlighted and displayed on the right. (B) Genomic peaks of P0 ATAC-seq (Aldiri et al., 2017), P0 Ikzf4 CUT&RUN replicate 1, P0 Ikzf4 CUT&RUN replicate 2, P0 IgG CUT&RUN at genomic tracks of Sox8, Sox9, Lhx2 and Nfi family members. Asterisks represent peaks called by MACS2. (C-D) RT-qPCR analysis of Müller specification gene expression from sorted GFP^+ve^ cells either 18 (C) or 72hours (D) after electroporation of P0 retinas with either GFP (n=5) or Ikzf4-IRES-GFP (n=5). (E-G’ ) Examples of GFP or Ikzf4-IRES-GFP electroporations at P0 and immunostained for Nfia/b 72 hours later. (H) Quantifications of GFP^+ve^Nfia/b^+ve^ cells 72 hours after electroporation in P0 retinas (GFP, n=6; Ikzf4, n=6). *p<0.05, ****p<0.0001. Statistics: Mann-Whitney test (C). Two-tailed unpaired t-test (G). RPL: Retinal progenitor layer. RQ: Relative quantitation. Scale bars: 10μm (D-F’).

To gain insights on how Ikzf4 might promote Müller glia development, we assessed genes enriched in the Müller glia cluster within P14 scRNA-seq data (Clark et al., 2019). Remarkably, we found Ikzf4 binding peaks in all of the top 8 genes enriched in the Müller glia cluster (Fig. S6A). However, as late RPCs and Müller glia have similar transcriptomes (Blackshaw et al., 2004; Roesch et al., 2008), these genes are also expressed in RPCs, complicating the interpretation of these results. Therefore, we focused our analysis on genes that are required for Müller glia development. We found Ikzf4 binding signal at genomic regions around *Sox8*, *Sox9, Lhx2*, and *Nfia/b/x* gene bodies, some of which were called as peaks by MACS2 (Fig. 5B) (Clark et al., 2019; de Melo et al., 2018; de Melo et al., 2016; Muto et al., 2009; Poche et al., 2008; Zibetti et al., 2019). In contrast, we did not find Ikzf4 binding at other intronic open chromatin regions in gene bodies of *Nfib*, *Nfix,* and *Nrl*, which are highly expressed at P0, showing the specificity of the identified Ikzf4 binding sites (Fig.S6B).

Next, we performed Transcription factor Occupancy prediction By Investigation of ATAC-seq Signal (TOBIAS) on open chromatin regions bound by Ikzf4 at E14 and P0 to assess transcription factor motif footprints (Bentsen et al., 2020). We found that, out of the 849 motifs analysed from JASPAR database (Fornes et al., 2020), motifs of transcription factors associated with Müller glia differentiation such as Lhx family, Sox family and Nfib were differentially enriched in P0 open chromatin regions with Ikzf4 binding compared to E14 (Supplementary file 4, Fig. S6C-D). We also found that, in the P0 retina, Ikzf4 binds to open chromatin regions that are active or poised enhancers, as suggested by the enrichment of H3K27ac and H3K4me3 histone marks, respectively (Creyghton et al., 2010; Orford et al., 2008), and depletion of the H3K27me3 histone mark (Aldiri et al., 2017; Cao et al., 2002) (Fig. S6E). Therefore, these data suggest that Ikzf4 binds to CREs important for Müller glia differentiation at late stages of retinal development.

We then assessed whether Ikzf4 regulates transcripts of genes involved in Müller glia development by electroporating P0 retinas with either CAG:GFP or CAG:Ikzf4-IRES-GFP and carried out RT-qPCR on sorted GFP+ cells 18 and 72 hours later. We found that Ikzf4 promotes expression of *Sox8* and *Sox9* at 18 hours post-electroporation. Although we found no change in *Lhx2*, *Rnf12*, *Nfia/b/x,* or *Ldb1* at 18 hours (Fig. 5C), their expression was increased 72hours after electroporation (Fig. 5D). To validate these RT-qPCR results, we electroporated P0 retinal explants with the same vectors and stained sections for Nfia/b 72hours later. As expected, we found a significant increase in the number of GFP^+ve^ cells stained for Nfia/b (Fig. 5C-H). These results suggest that Ikzf4 promotes Müller glia production by binding to and upregulating genes involved in glial cell specification, including the Nfi temporal identity factors.

### Ikzf4 upregulates Hes1 by binding to a consensus Ikaros binding site in the promoter

In our CUT&RUN data, we noticed that the *Hes1* and *Hes5* promoters are highly bound by Ikzf4, both at early and late stages of retinogenesis. It was previously shown that Hes1 expression oscillates during progenitor proliferation (Shimojo et al., 2008). Interestingly, Hes1 expression decreases when the RPCs exit the cell cycle and is repressed in cells fated to become neurons, whereas it is maintained in cells fated to become glia (Furukawa et al., 2000; Imayoshi et al., 2013). To study the dynamics of Ikzf4 binding at the *Hes1* and *Hes5* promoter, we co-electroporated reporter constructs pHes1-dsRed or pHes5-dsRed together with either CAG:GFP or CAG:Ikzf4-IRES-GFP in P0 retinas. We tracked the electroporation patches over time to assess the dynamics of dsRed expression, which reports activity of the promoters. We observed that Ikzf4 promotes the activity of the Hes1 promoter compared to the control GFP at 48 and 72 hours (Fig. S7A-D’’). Even 6 days after electroporation, Ikzf4-electroporated retinas maintained high expression of dsRed, whereas expression of dsRed was reduced in the control GFP condition (Fig. S7E-F’’). In contrast, we did not observe similar dynamics in dsRed expression for the Hes5 promoter at 48hours or 6 days after Ikzf4 electroporation compared to the GFP control (Fig. S7G-J’’). Of note, Ikzf4-mediated upregulation of the *Hes1* promoter activity appears to be specific to retinal cells, as Ikzf4 had no effect on pHes1-dsRed activity in HEK293 cells (Fig. S7K-L’’). We next assessed whether the increase in *Hes1* promoter activity leads to an increase in Hes1 protein expression. To test this, we electroporated P0 retinas with either CAG:GFP or CAG:Ikzf4-IRES-GFP and analyzed the number of GFP^+ve^ cells staining for Hes1 in retinal sections 44 hours later. As expected, we observed an increase in the number of GFP^+ve^Hes1^+ve^ cells after Ikzf4 expression (Fig. 6A-C). Together, these results indicate that Ikzf4 activates the Hes1 promoter, which leads to elevation of Hes1 protein expression in a larger proportion of cells.

**Figure 6:**
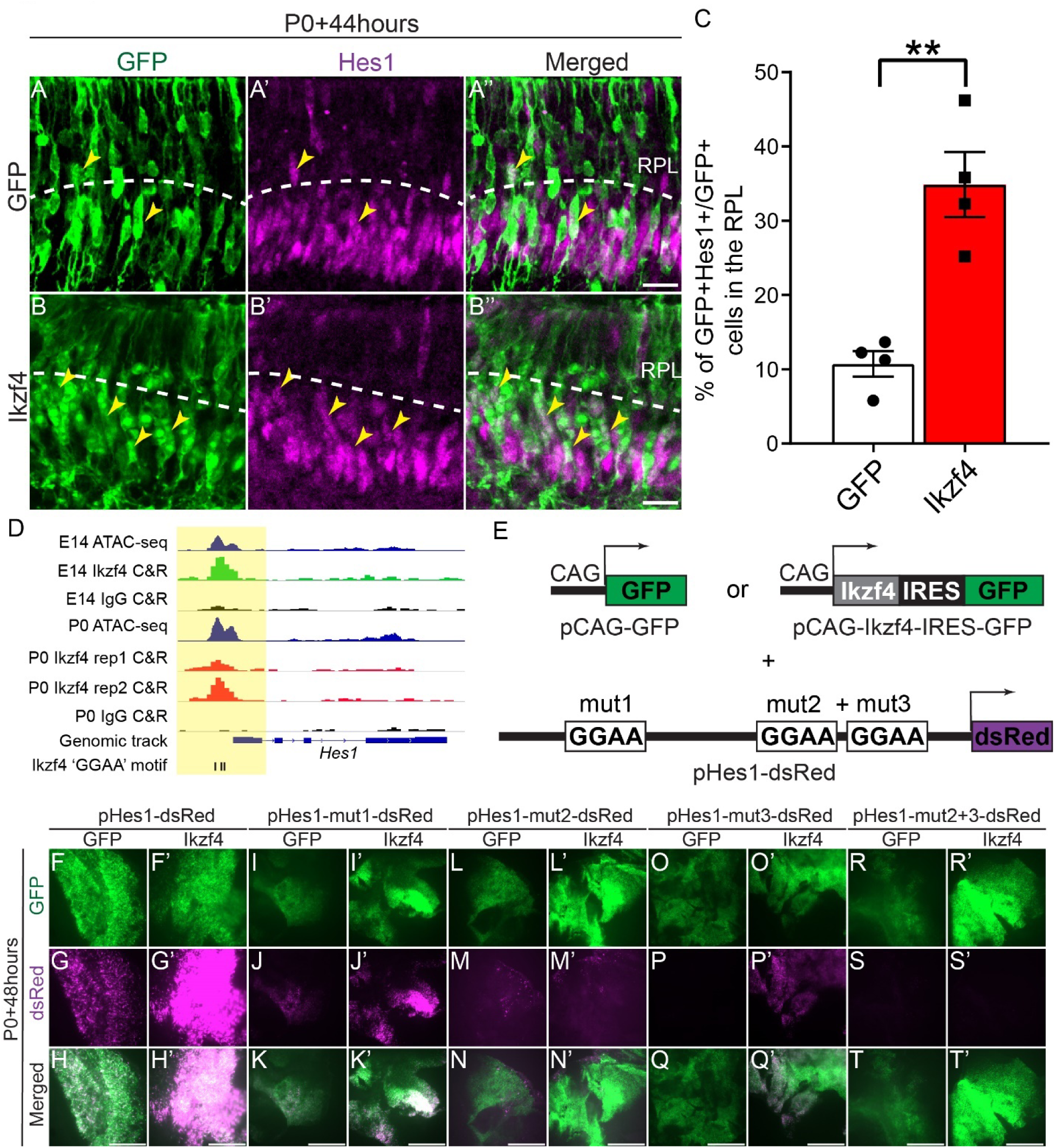
Ikzf4 binds to the Hes1 promoter and upregulates Hes1 expression. (A) Examples of GFP or Ikzf4-IRES-GFP-electroporated P0 retinas immunostained with Hes1 44 hours later. Yellow arrows indicate GFP^+ve^Hes1^+ve^ cells in the RPL. (C) Quantification of GFP^+ve^Hes1^+ve^ cells in the RPL at P0+44 hours after electroporation (GFP, n=4; Ikzf4, n=4). (D) Genomic peaks of E14 or P0 ATAC-seq (blue) (Aldiri et al., 2017), E14 Ikzf4 CUT&RUN (green), E14 and P0 IgG CUT&RUN (black), P0 Ikzf4 CUT&RUN replicate 1 (red), P0 Ikzf4 CUT&RUN replicate 2 (red) at Hes1 promoter. Ikzf4 ‘GGAA ’motifs are denoted as black bars. Yellow highlighted area is 500bp around the promoter region of Hes1. (E) Schematic representation of experiment shown in (F-T’). Retinal explants were co-electroporated with either pCAG-GFP or pCAG-Ikzf4-IRES-GFP along with vectors expressing dsRed under the control of either WT Hes promoter (pHes1-dsRed), Hes1 promoter with mutation at first ‘GGAA ’ site (mut1), second ‘GGAA ’site (mut2), third ‘GGAA ’(mut3) site and second and third GGAA mutations together (mut2+mut3). (F-T’) Photomicrographs of retinal flatmounts showing increase in dsRed expression when Ikzf4-IRES-GFP is co-electroporated with either pHes1-dsRed (F’-H’), pHes1-mut1-dsRed (I’-K’), pHes1-mut2-dsRed (L’-N’), pHes1-mut3-dsRed (O’- Q’) but not with pHes1-mut2+3-dsRed (R’-T’) compared to the control GFP (F-G, I-K, L-N, O-Q, R-T). Statistics: Two tailed unpaired t-test (C). **p<0.01. RPL: Retinal progenitor layer. Scale bars: 10μm (A-B”), 250μm (F-T’).

Finally, we sought to identify the Ikzf4 binding sites required for the regulation of *Hes1*. When we analyzed the *Hes1* promoter region containing the Ikzf4 peaks, we found three ‘GGAA’ Ikzf binding motifs (Fig. 6D). To study the functional requirement of these motifs, we mutated each one and assessed how this affected the ability of Ikzf4 to regulate the activity of the Hes1 promoter. We co-electroporated CAG:GFP or CAG:Ikzf4-IRES-GFP with either the WT or mutated *Hes1* promoter constructs (Fig. 6E). When we mutated binding site 1 (mut1), 2 (mut2) or 3 (mut3) and electroporated with CAG:GFP, we observed a reduction in dsRed signal compared to the WT construct (Fig. 6F-Q’), confirming the importance of these sites for the activity of the *Hes1* promoter. However, co-expression of Ikzf4 was sufficient to promote dsRed expression, albeit less so for mut2 (Fig. 6F’-Q’). This suggests that none of these sites alone is required for Ikzf4 binding and activation of the Hes1 promoter (Fig. 6F-Q’). Interestingly, however, we found that when we mutated sites 2 and 3 together (mut2+mut3), Ikzf4 was no longer able to activate the Hes1 promoter (Fig. 6R-T’). Taken together, these results show that Ikzf4 binds at two ‘GGAA’ sites that are required for the sustained expression of Hes1 in post-mitotic cells.

## DISCUSSION

Neural diversity in the CNS is generated by a combination of spatial and temporal factors working in concert to establish the vast repertoire of neurons and glia. While factors regulating fate decisions upon cell cycle exit have been extensively studied, much less is known about how neural progenitors alter their developmental competence over time. In this study, we show that Ikzf4 cooperates with the previously identified early temporal identity factor Ikzf1 to control the production of both early-born cone photoreceptors and late-born Müller glia. Mechanistically, we report that, during early stages of development, Ikzf1 and Ikzf4 are both expressed in RPCs and generally bind the same target genes involved in cone specification, but at different CREs. At late stages, Ikzf1 is no longer expressed, but Ikzf4 expression is maintained and switches targets to bind genes involved in Müller glia development, including Sox8, Sox9, Lhx2, Nfi, and the Notch signaling effector Hes1. Ikzf4 induces sustained expression of Hes1 during differentiation via two ‘GGAA’ Ikzf binding sites in the promoter. Taken together, this study identifies functional cooperation between Ikaros family proteins and dynamic regulation of target gene selection as key mechanisms underlying temporal patterning in neural progenitors (Fig. 7).

**Figure 7:**
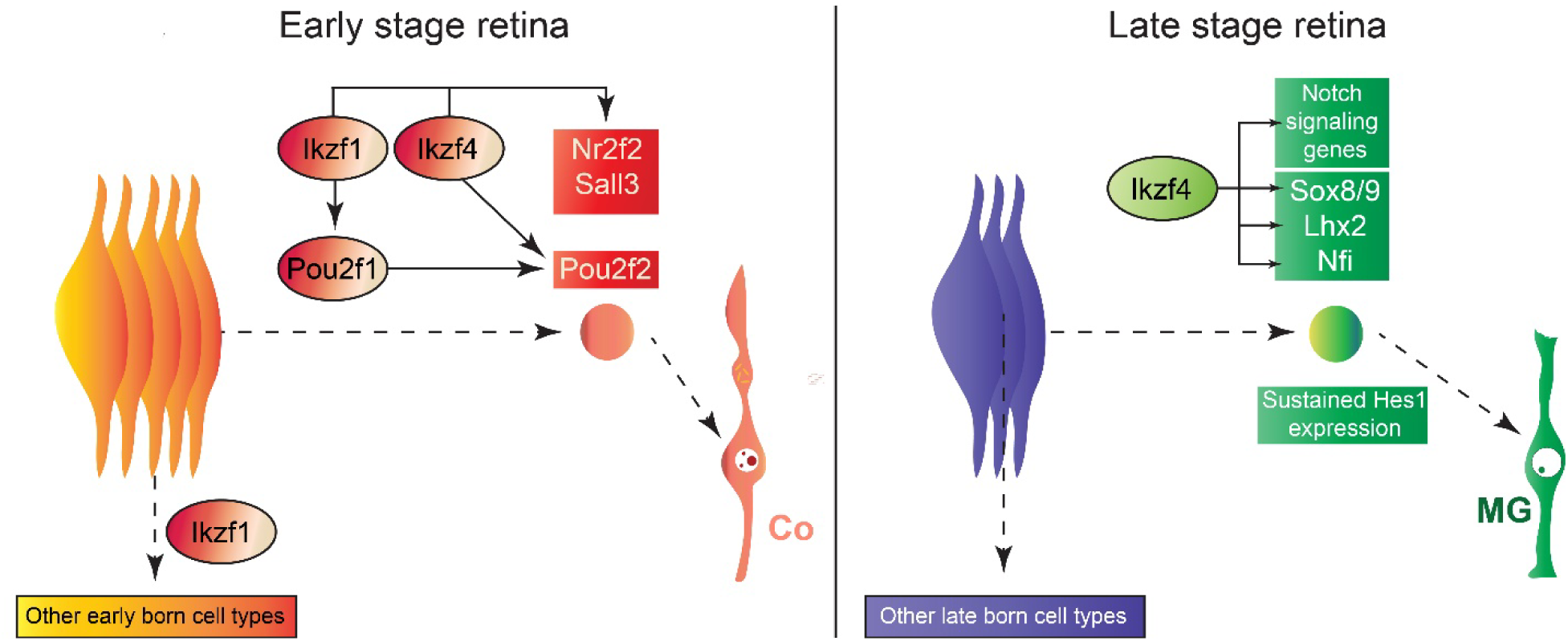
Model of temporal patterning during mouse retinogenesis. In early RPCs, Ikzf4 functions redundantly with Ikzf1 to confer competence for cone (Co) production, whereas Ikzf1 regulates competence for production of other early-born cell types. In late RPCs, Ikzf4 acts as a classical fate determinant by binding and upregulating expression of Müller specification genes as well as instigating the maintenance of Notch signaling to ensure the Müller glia (MG) fate commitment.

### Ikzf1 and Ikzf4 in the control of cone development

Single knockout retinas of Ikzf1 or Ikzf4 have normal number of cones (Fig. 3C) (Elliott et al., 2008), but here we report that double knockouts of Ikzf1 and Ikzf4 have fewer cones, suggesting genetic redundancy for cone production. Although we find that Ikzf1 and Ikzf4 bind different CREs, we also report general co-binding at the same genes, such that inactivation of only one factor likely leads to compensation by the other factor in cone gene regulation. It remains unclear, however, whether Ikzf1 switches its binding profile upon loss of Ikzf4 and vice versa. A comprehensive analysis of genomic binding of Ikzf family members in KO retinas lacking other Ikzf factors will be required to explore this possibility.

Our data indicate that Ikzf4 acts upstream of *Pou2f2* to initiate cone genesis at the appropriate developmental time. We observe an increase in Rxrg^+ve^ cells when Ikzf4 is expressed in late RPCs, suggesting that Ikzf4 is sufficient to reopen a window of competence for cone genesis that is normally lost at this stage. However, Ikzf4 expression in late RPCs does not lead to the production of mature cones, unlike Pou2f1 (Javed et al., 2020). This suggests that Ikzf4 is able to promote cone genesis, but likely requires additional factors that are missing at late stages of retinogenesis to induce their full maturation. Ikzf4 also promotes early cell cycle exit, which is uncharacteristic of temporal identity factors like Ikzf1, Pou2f1 and Casz1, but not Foxn4 (Elliott et al., 2008; Javed et al., 2020; Li et al., 2004; Mattar et al., 2015). Altogether, our results show that Ikzf4 is an important part of a gene regulatory network that confers RPCs with the competence to generate cones.

### Ikzf4 in the regulation of Müller glia production

Our data indicate that Ikzf4 switches transcriptional targets at late stages of retinogenesis to control Müller glia production. One possible mechanism to explain this switch is that Ikzf4 is co-expressed with Ikzf1 during early stages of retinal development, but not at late stages. The absence of Ikzf1 at late stages of development may redirect Ikzf4 from cone genes to glial genes, although such activity would have to be indirect since Ikzf1 and Ikzf4 have divergent genome occupancy. The current model of gliogenesis in the retina proposes that Nfia/b/x confer late-stage temporal identity to RPCs to generate Müller glia and bipolar cells (Clark et al., 2019; Lyu et al., 2021). In addition to Nfia/b/x, Lhx2 interacts with Rnf12 in late RPCs to promote gliogenesis by activating the expression of *Sox8/9* and Notch target genes in postmitotic precursors destined to become Müller glia (de Melo et al., 2018; de Melo et al., 2016; Jadhav et al., 2006; Muto et al., 2009; Nelson et al., 2011; Poche et al., 2008; Zhu et al., 2013; Zibetti et al., 2019). Lhx2 also dynamically alters its DNA binding profile at early and late stages of retinal development to switch from promoting neurogenesis to gliogenesis (Zibetti et al., 2019), similar to what we observe here with Ikzf4. So how does Ikzf4 fit in the above proposed model of gliogenesis? As Lhx2 and Nfia/b/x cKOs show no significant change in *Ikzf4* transcript levels (Clark et al., 2019; de Melo et al., 2016), and our data show that Ikzf4 binds to and induces expression of Lhx2 and Nfi, we propose that Ikzf4 may be upstream of these factors in the gliogenic gene regulatory network. Given the weak expression levels of *Ikzf4* and the low sensitivity of the RNA-seq analysis, however, it is possible that previous studies failed to detect changes in *Ikzf4* expression in the Lhx2 and Nfi cKOs. In any case, our data suggests a model wherein Ikzf4 favours the emergence of Sox8/9^+ve^ precursors with sustained Notch signaling, as observed with the increase in Hes1^+ve^ cells after overexpression of Ikzf4 in RPCs, which then go on to become Müller glia.

We report that Ikzf4 binds to regions close to many Notch signaling gene bodies, including multiple sites at the promoter of Hes1. As the Hes1 promoter is active in RPCs and differentiated Müller glia (Matsuda and Cepko, 2007), this suggests that Ikzf4 might be involved in maintaining *Hes1* expression in mature glia, although this remains to be tested directly. Using CUT&RUN sequencing, we observe a change in genomic binding profiles of Ikzf4 from early to late stages of retinal development, such as binding to some Notch signaling genes that are exclusively observed at P0, for example *Notch1* and *Rbpj* (Fig. 5A). This suggests that Ikzf4 regulation of Notch signaling may be more prominent during late stages of retinogenesis, but this remains to be tested. Interestingly, it was recently reported that *Hes1* deletion in early RPCs leads to a reduction in cone numbers and impaired Müller glia morphology (Bosze et al., 2020), two cell fates regulated by Ikzf4, supporting the idea that *Hes1* is a key target of Ikzf4, both during early and late retinogenesis.

### Broader role of Ikzf4 in cell fate specification

Previous studies have elucidated the role of Ikzf4 in the immune system. Most notably, Ikzf4 function in T-cell differentiation varies considerably depending on CD4^+^ T-cell subtype, highlighting the dynamic role of Ikzf4 function based on the cellular context (Liu et al., 2014; Pan et al., 2009; Powell et al., 2019; Read et al., 2017; Rieder et al., 2015; Sekiya et al., 2015; Sharma et al., 2013), similar to what we report here. Interestingly, widespread expression of Ikzf4 has been reported in different organs, including the CNS (Perdomo et al., 2000). One study has detailed the role of Ikzf4 in regulating expression of PSD-95 in adult cochlear afferent neurons (Bao et al., 2004), but the function of Ikzf4 in the developing CNS remains largely elusive. The findings reported here suggest that Ikzf4 may have widespread regulatory functions in the CNS to control cell fate specification. Future studies on conditional knockouts of Ikzf4 alone or in combination with other Ikzf genes will be interesting to address this question.

## MATERIALS AND METHODS

### Animals

All experiments were done in accordance with the Canadian Council on Animal guidelines. Ikzf1 (Wang et al., 1996) and Ikzf4 (International Mouse Phenotyping Consortium and RIKEN Bioresource **<u>RRID: IMSR_RBRC06808</u>**) knockout mice were raised in the C57BL/6J background (Mus musculus). All other mouse experiments were performed on WT CD1 mice (Mus musculus, Charles River Laboratories).

### Retroviral constructs preparation and retinal explant culture

Retroviruses were designed, produced and concentrated as previously described (Cayouette et al., 2003). Retinal explants were cultured as previously outlined (Cayouette et al., 2001). Retroviral infections of retinal explants and analyses of the retroviral clones were done as previously stated (Javed et al., 2020). Eyes were harvested from the mice 14 days or more after electroporation as required for the experiment and processed for immunostaining.

### Plasmid and mutation cloning

Ikzf1 and Ikzf4 cDNA were cloned into a pCIG2-IRES-GFP and pCLE-venus vector using restriction sites previously outlined (Gaiano et al., 2000; Hand et al., 2005). pHes1-dsRed vector was mutated with Infusion HD Cloning Plus kit from Takara using primers listed in Table S1 (Matsuda and Cepko, 2007).

### *In vivo* electroporation

P0 or P1 eyes were injected with DNA plasmid at 3ug/ul concentration containing 0.5% fast green and electroporated as previously described (de Melo and Blackshaw, 2011).

### RNA isolation and Quantitative PCR

RNA extraction and qPCR quantitation were performed as previously described (Javed et al., 2020). Primers used are listed in Table S1 (Batsché et al., 2005; Ouimette et al., 2010).

### RNA-seq preparation

P0 CD1 retinas were dissociated using Accutase and transfected using the Amaxa Neural Stem Cell Kit according to the manufacturer’s protocol. Cells were seeded into 6-well plates coated with PLL/ Laminin, and cultured as previously described (Gomes et al., 2011). After 9 hr, cells were harvested and dissociated with Accutase followed by FAC-sorting for GFP. Control GFP condition contained 4 x10^5^ cells per biological replicate (n=4) and Ikzf1 condition contained 3 x 10^5^ cells per biological replicate (n=4). Cells were sorted directly into lysis buffer and purified using the RNeasy micro kit from Qiagen. Samples quantity (RNA Integrity Number = RIN) and quality were assessed using the Agilent RNA 6000 Pico Kit (Agilent #5067-1513) on the Bioanalyzer 2100. rRNA was depleted using Ribo-Zero™ Magnetic Gold Kit for rRNA depletion (Human/Mouse/Rat) (Epicentre for Illumina #MRZG12324) according to manufacturer’s guidelines. Library preparations were done using SMARTer® Stranded RNA-Seq Kit (Clontech Laboratories, Inc. #634836 or #634837) according to manufacturer’s guidelines. Libraries were diluted and pooled equimolar and then sequenced in pair end 50 cycles (PE50) on a v4 flowcell (Illumina - HiSeq PE Cluster Kit v4 cBot - PE-401-4001) of the Illumina HiSeq 2500 System.

### Tissue collection and immunofluorescence

Age of mouse embryos was calculated from pregnant females with the day of vaginal plug considered as day 0 (E0) and collected at E11, E14, E15, E16, E17, P0, P2, and P7 for spatiotemporal analyses. For Ikzf1 and Ikzf4 immunostaining, the retinas were dissected and fixed for 2mins in 4%PFA/PBS followed by immersion in 20% Sucrose/PBS for 1hour. Retinas were then embedded in OCT, frozen in liquid nitrogen, sectioned at 25μm using a cryostat and immunostained on the same day. For all other antibodies, the decapitated heads from embryos or eyes from postnatal pups were fixed for 15mins in 4%PFA/PBS and immersed in 20% Sucrose/PBS for 2 hours.

Immunofluorescence was performed as previously described (Javed et al., 2020). List of primary antibodies can be found in Table S2 including DHSB antibodies (Venkataraman et al., 2018).

### EdU labelling assay

30μM of EdU was added to the culture medium for 2hours before collection and fixation. Click-iT^TM^ EdU Alexa Fluor^TM^ 647 was used to label cells that incorporated EdU.

### Statistical and Quantitative analyses

Statistical tests were performed for each experiment in this study as indicated in the Figure legends. All quantifications in the bar graphs of this study are represented as mean ± standard error of the mean (s.e.m.) whereas n number and individual values on the graphs represent biological replicates. Statistics for the retroviral clonal analyses were performed as previously outlined (Pounds and Dyer, 2008). Retinal explants containing disorganised layers and poor immunostainings were discarded and analysis was limited to the organised regions of the retinal explants. All experiments were repeated at least three times.

Ikzf4 knockouts were analysed as follows: Two sections of P10 central retinas oriented temporo-nasally were examined with quantifications of 200μm (Pax6), 400μm (Otx2, Sox2, and Lhx2 staining in the INL, Rxrg in the ONL) and 800μm (Brn3a and Lim-1) length of the imaged section. The investigator was blinded to the genotype of animals. An ImageJ analysis macro was written to count cell automatically in a section. Analyse particles was used after defining a region of interest and setting threshold for each antibody to auto-count cell numbers of Pax6^+ve^, Otx2^+ve^, Sox2^+ve^, Lhx2^+ve^ and Brn3a^+ve^ cells whereas Rxrg^+ve^ and Lim-1^+ve^ were manually counted. Ikzf1/4 double knockouts were analysed as follows: Three sections of E15 central retinas in embryonic heads oriented dorso-ventrally were analysed with quantification of 200μm for Rxrg^+ve^ cells and 400μm for Brn3a^+ve^ cells. Rxrg^+ve^ cell were manually counted due to background signal, whereas Brn3a^+ve^ cells were counted using the ImageJ analysis macro as described above.

All cell quantifications were performed by blinding the investigators to the genotype of the animals.

### CUT&RUN assays

CUT&RUN was performed as previously described (Skene et al., 2018), with a few added modifications. For Ikzf1 CUT&RUN, CAG-Ikzf1 electroporated P0 retinal explants were cultured and 50,000 GFP+ve cells were sorted directly into the buffer with Concanavalin-A beads. For Ikzf4 CUT&RUN, E14 and P0 retinas were dissected, and 1,500,000 cells were used for each stage. The entire procedure was done in 200μl PCR tubes. 0.01% digitonin concentration was used and pAG-MNase digestion was performed for 30min on ice.

Libraries were prepared with the KAPA DNA HyperPrep Kit (Roche 07962363001 - KK8504). This protocol includes an End-Repair/A-tailing step and an adapter ligation step followed by a PCR amplification (enrichment) of ligated fragments. The adapters used for ligation were IDT for Illumina TruSeq UD Indexes (Illumina - 20022371). The final enriched product (library, after PCR) was purified using KAPA purification beads (Roche 07983298001 - KK8002) and a dual-SPRI size selection was performed (with KAPA beads) to select fragments between 180-500 bp. Libraries were then quantified using a Nanodrop microvolume spectrophotometer (ng/μl) and quality was assessed using the Agilent High Sensitivity DNA Kit (Agilent -5067-4626) on a Bioanalyzer 2100. The libraries were then quantified by q-PCR to obtain their nanomolar (nM) concentration. Libraries were diluted, pooled equimolar and sequenced in pair end 50 cycles (PE50) on a S1 flowcell (Illumina - 20012863) of the Illumina NovaSeq 6000 System.

### Bioinformatics analyses

scRNA-seq analyses of previously published datasets was performed as follows. Fastq raw reads were aligned and counted to generate matrices for each individual stage from Mouse retina atlas (GEO: GSE118614) (Clark et al., 2019) and Human fetal retina atlas (GEO: GSE116106, GSE122970, GSE138002) (Lu et al., 2020) using Cellranger 4.0 (10x Genomics). Velocyto (La Manno et al., 2018) with run10x function was used on cellranger output folders to generate .loom files for each individual stages. Seurat (Butler et al., 2018) was used to analyse the mouse and human retina. loom files and subset RPC clusters for Ikzf4/IKZF4 expression analyses using markers previously described (Clark et al., 2019; Lu et al., 2020). Scanpy (Wolf et al., 2018) was used to analyse the same .loom files and generate Ikzf4/IKZF4 expression UMAPs along with cell type markers (Fig. S2).

RNA-seq fastq raw reads were analysed using the salmon quantification package (Patro et al., 2017). Differentially gene expression analyses between CAG-GFP and CAG-Ikzf1 was performed using DESeq2 on Salmon quant output files (Love et al., 2014). Microsoft Excel was used to compare differentially expressed gene lists with other datasets manually.

CUT&RUN analyses were performed by aligning the raw fastq reads with the mouse mm9 genome using default parameters of bowtie2 on Galaxy platform (Afgan et al., 2016; Langmead and Salzberg, 2012). Bam files generated for Ikzf4 and IgG CUT&RUN at each stage were used to call peaks using 0.5 FDR parameter on MACS2 (Feng et al., 2012). Bedtools was used to find overlapping peaks between the two stage (Quinlan and Hall, 2010). Bigwig files were generated using deeptools2 (Ramirez et al., 2016). Integrative Genomics Viewer (IGV) was used to visualize bigwig files.

TOBIAS footprinting analysis was performed by first intersecting ATAC-seq peaks at E14 and P0 from previously published datasets (Aldiri et al., 2017), with Ikzf4 E14 and P0 regions using bedtools (Bentsen et al., 2020; Quinlan and Hall, 2010). TOBIAS ATACorrect and ScoreBigwig was performed on ATAC-seq E14 and P0 .bam files separately to generate corrected ATAC-seq bigwig files. The two E14 and P0 regions were merged using bedtools and BINDetect was used to estimate differentially bound motifs based on scores, sequence and motifs. JASPAR non-redundant vertebrate motifs package was used for motif annotation on the mm9 genome (Fornes et al., 2020).

Volcano plot was generated using the EnhancedVolcanoplot github package (https://github.com/kevinblighe/EnhancedVolcano), whereas Upset plots were generated using Upsetplot shiny web tool (Conway et al., 2017).

Information on each software version is listed in Table S2.

### Data availability

E14 IgG CUT&RUN raw reads are available on previously published GEO accession number: GSE156756 (Brodie-Kommit et al., 2021). Processed RNA-seq, MACS2 peaks and bigwig files are available on GEO accession number: GSE189590.

## Supporting information

Excel spreadsheet containing the complete list of differential enriched motifs between Ikzf4 E14 and Ikzf4 P0 open chromatin regions using TOBIAS BIND

Excel spreadsheet showing GREAT analysis of Ikzf4 E14 exclusive, P0 exclusive and Common binding target gene and peak location

Excel spreadsheet describing GOnet GO term annotation of the genes found in Supplementary file 1

Excel spreadsheets containing the genes up/downregulated and bound by Ikzf1 compared to genes bound by Ikzf4 at E14 and P0 after GREAT analysis

## ACKNOWLEDGEMENTS

We thank Christine Jolicoeur, Jessica Barthe, Androne Constantin, Odile Neyret- Djossou, Éric Massicotte and Julie Lord for animal and technical assistance. We thank Seth Blackshaw for critical comments on the manuscript. We thank Aurelie Huang-Sung and Dr. Nicole Francis for providing the pAG-MNase for the CUT&RUN experiments. We thank Benoit Boulan for writing the ImageJ macro. We also thank all members of the Cayouette lab for their comments and support. This project was funded by grants from the Canadian Institutes of Health Research (FDN-159936) and Fighting Blindness Canada to M.C. A.J. was supported by a Ph.D. scholarship from the Fonds de recherche du Québec – Santé. M.C. is an Emeritus Scholar from Fonds de recherche du Québec – Santé and holds the Gaëtane and Rolland Pillenière Chair in Retinal Biology from the Montreal Clinical Research Institute Foundation.

## AUTHOR CONTRIBUTIONS

Conceptualization, A.J. and M.C.; Investigation, A.J., P.M., A.C.; Writing – Original Draft, A.J.; Writing – Review & Editing, P.M., M.C., and M.C.; Resources, M.C.; Supervision, M.C.; Funding Acquisition, M.C.

