## Supplemental Figures and Tables for "Ikaros family proteins regulate developmental windows in the mouse retina through convergent and divergent transcriptional programs"

### SUPPLEMENTARY FIGURES:

Figure S1, Javed et al.

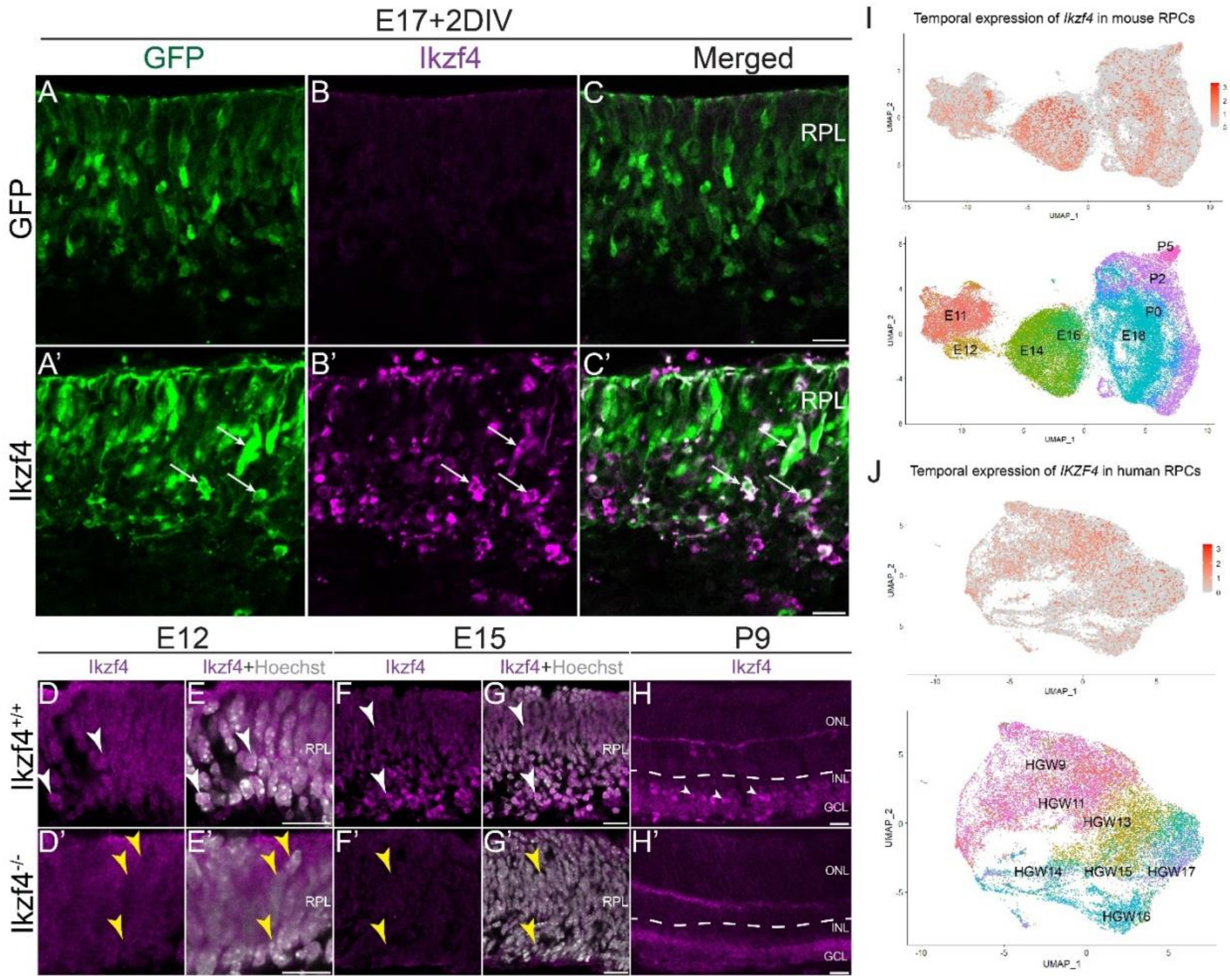


**Figure S1: Specificity of Ikzf4 antibody and scRNA-seq re-analysis of human and mouse RPCs.**

(A-C’) Examples of electroporation of either GFP (A-C) or Ikzf4-IRES-GFP (A’-C’) in E17 retinas and immunostained with Ikzf4 antibody 2 days later. White arrows indicate GFP$^{+ve}$Ikzf4$^{+ve}$ cells. (D-H’) Validation of the Ikzf4 antibody in Ikzf4$^{+/+}$ retinas (D-H) compared to Ikzf4$^{-/-}$ retinas (D’-H’) at E12 (D-E’), E15 (F-G’) and P9 (H-H’). White arrows show Ikzf4$^{+ve}$ cells with nuclear immunostaining and yellow arrow indicate absence of nuclear immunostaining. (I-J) Re-analysis of previously published single cell RNA-seq datasets from mouse (Clark et al. 2019) and human fetal (Lu et al. 2020) retinas. RPL: Retinal progenitor layer. ONL: Outer nuclear layer. INL: Inner nuclear layer. GCL: Ganglion cell layer. RPC: Retinal Progenitor Cell. HGW: Human Gestational Week. Scale bars: 10μm (A-H’).

Figure S2, Javed et al.

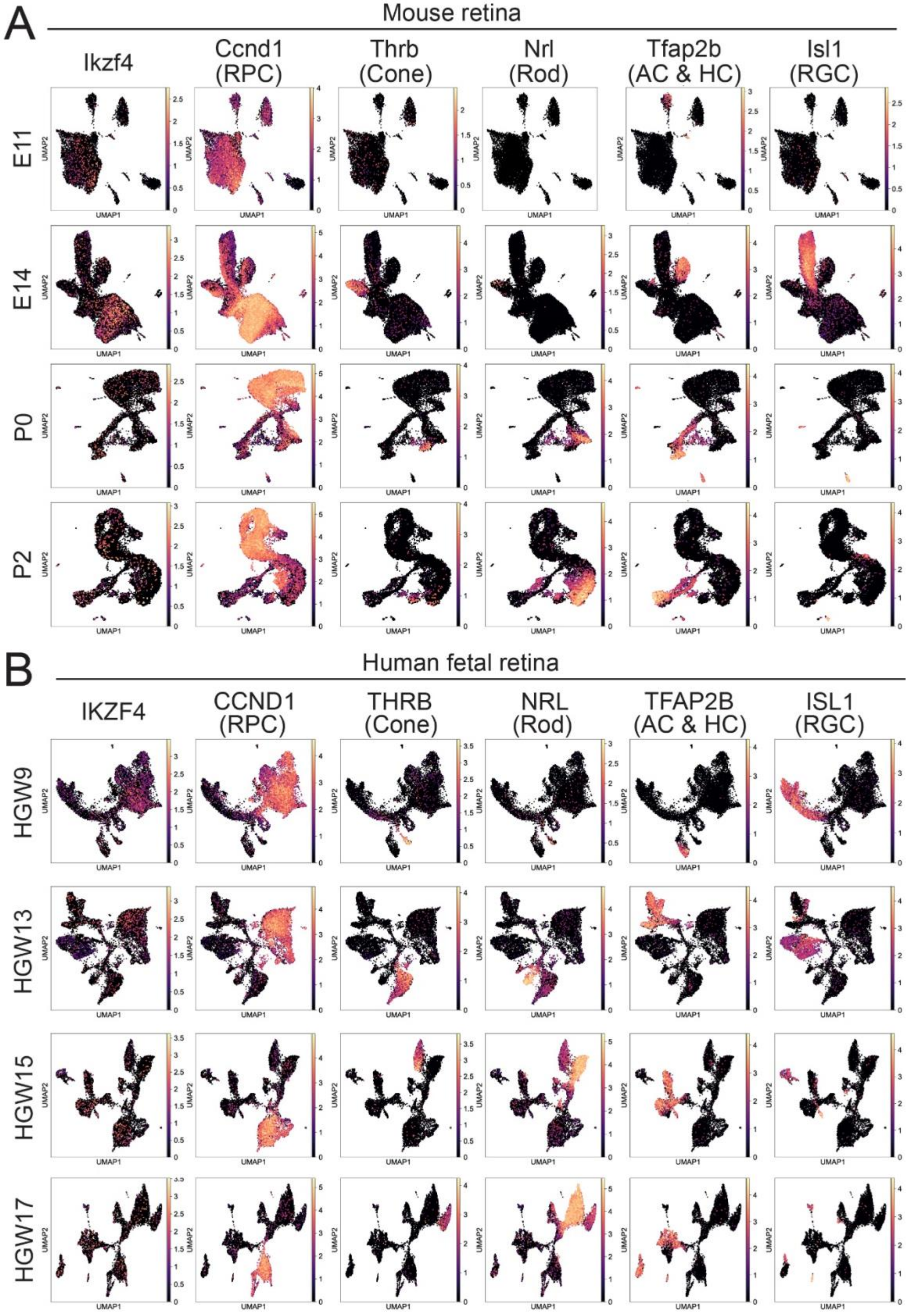

**Figure S2: *Ikzf4* mRNA is expressed in some differentiated cell types in mouse and human retina.**

(A) *Ikzf4* mRNA expression during mouse retinogenesis in RPCs (*Ccnd1*), cones (*Thrb*), rods (*Nrl*), amacrines and horizontal cells (*Tfap2b*) and RGCs (*Isl1*). (B) *IKZF4* mRNA expression during early to late human fetal retinogenesis in RPCs (*CCND1*), cones (*THRB*), rods (*NRL*), amacrines and horizontal cells (*TFAP2B*) and RGCs (*ISL1*).

Figure S3, Javed et al.

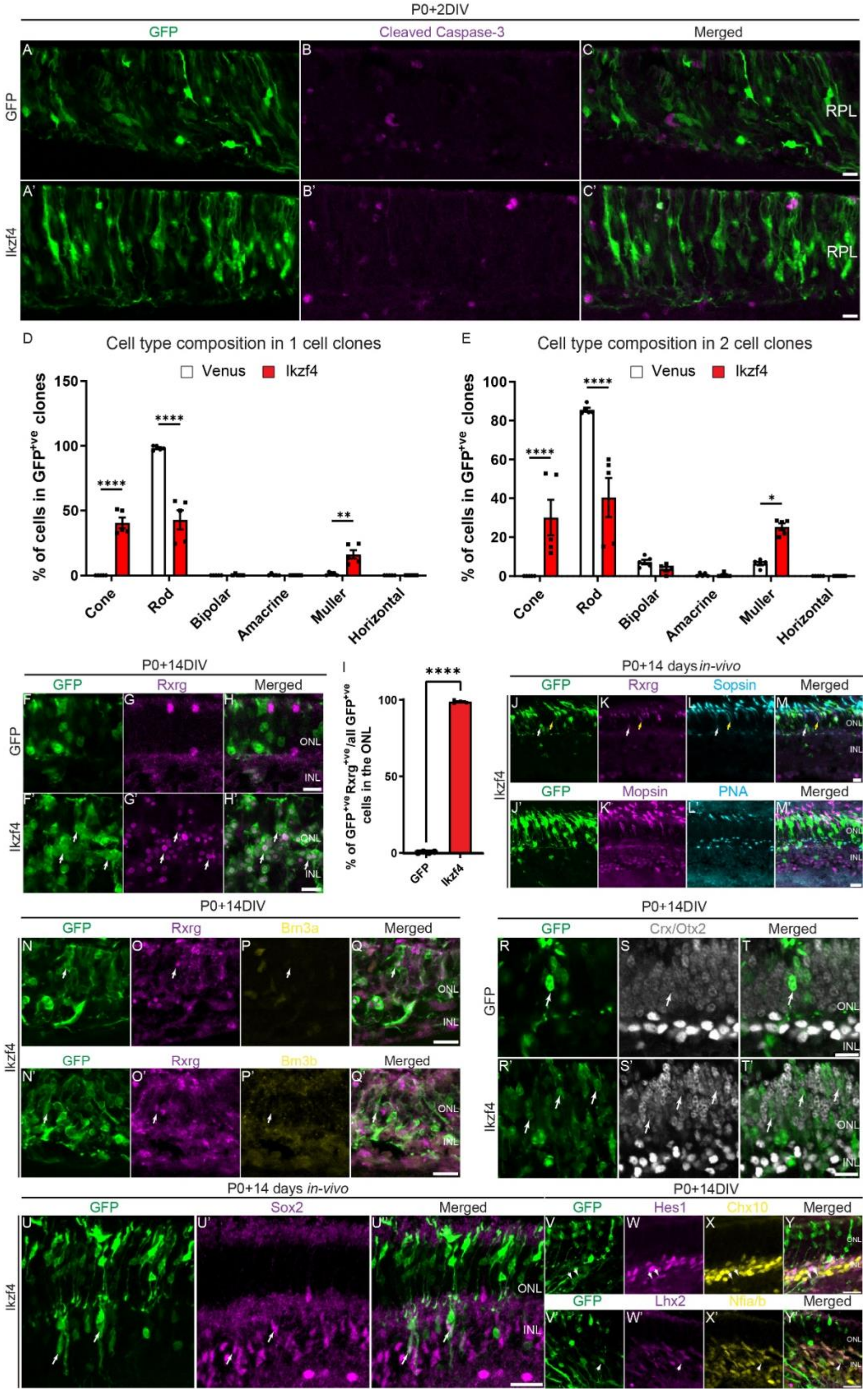

**Figure S3: Ikzf4 promotes cones and Müller glia from late RPCs by inducing early cell cycle exit rather than apoptosis.**

(A-C') Examples of retinal explants electroporated at P0 with either GFP (A-C) or Ikzf4-IRES-GFP (A'-C') and immunostained with Cleaved Caspase-3 (B-B') 2 days later. (D-E) Retroviral clonal analysis of Venus or Ikzf4 as shown in (Fig. 2A-G) focusing on cell type composition of 1 cell (D) or 2 cell (E) clones. (F-H') Examples of retinal explants electroporated at P0 with either GFP (F-H) or Ikzf4-IRES-GFP (F'-H') and immunostained with Rxrg (G-G') 14 days later. White arrows denote electroporated cells expressing Rxrg. (I) Quantification of retinal explants electroporated with either GFP (n=4) or Ikzf4-IRES-GFP (n=4). (J-M') Examples of in-vivo electroporations at P0 with Ikzf4-IRES-GFP (J-M') and immunostained 14 days later with Rxrg (K), S-opsin (L), M-opsin (K') or PNA (L'). White arrows denote GFP$^{+ve}$Rxrg$^{+ve}$S-opsin$^{-ve}$ cells whereas yellow arrow represent endogenous non-electroporated Rxrg$^{+ve}$S-opsin$^{+ve}$ cones. (N-Q') Examples of Ikzf4-IRES-GFP (N-Q') electroporations of retinal explants at P0 and co-immunostained 14 days later with Rxrg (O-O') with either Brn3a (P) or Brn3b (P'). (R-T') Examples of retinal explants electroporated at P0 with either GFP (R-T) or Ikzf4-IRES-GFP (R'-T') and immunostained with Crx/Otx2 (S-S') after 14 days of culture. (U-U'') Examples of in-vivo electroporation of P0 retinas with Ikzf4-IRES-GFP (U-U'') and immunostained with Sox2 (U') 14 days later. (V-Y') Examples of retinal explants electroporated at P0 with Ikzf4-IRES-GFP (V-Y') and immunostained with Hes1 (W), Chx10 (X), Lhx2 (W') or Nfia/b (X'). White arrows represent either Hes1$^{+ve}$Chx10$^{-ve}$ cells (V-Y) or Lhx2$^{+ve}$Nfia/b$^{+ve}$ cells (V'-Y'). *p<0.05, **p<0.01, ****p<0.0001. Statistics: Two tailed unpaired t-test (D-E, I). ONL: Outer nuclear layer. INL: Inner nuclear layer. RPL: Retinal progenitor layer. Scale bars: 10µm (A-C', F-H', J-T', V-Y'), 20µm (U-U'').

Figure S4, Javed et al.

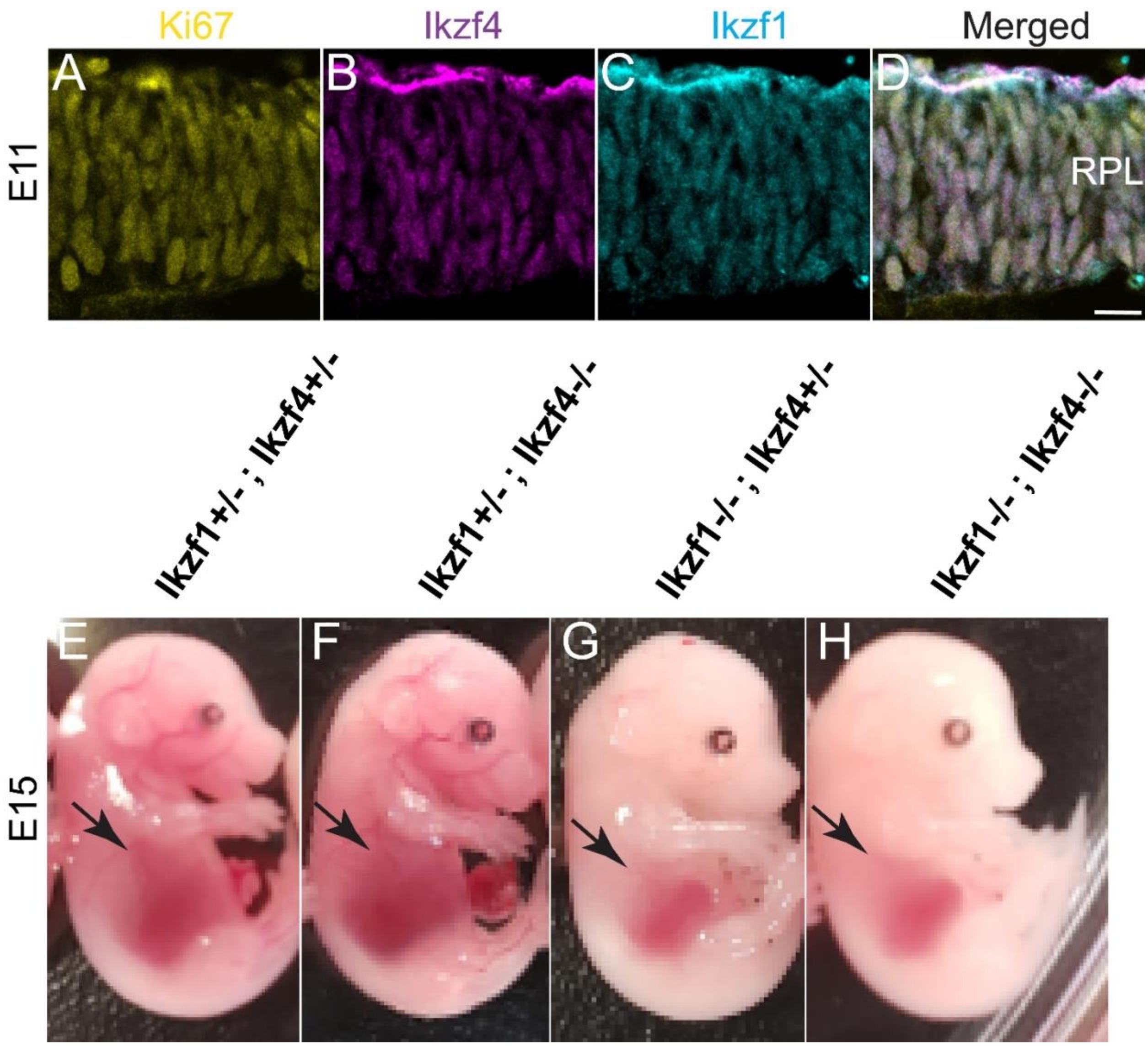


**Figure S4: Ikzf1 and Ikzf4 are expressed in the same cells during early retinogenesis and redundantly required for fetal liver size.**
(A-D) Co-immunostaining of Ki67 (A), Ikzf4 (B), Ikzf1 (C) in E11 retinas. (E-H) Examples of E15 embryos with genotypes, Ikzf1$^{+/-}$;Ikzf4$^{+/-}$ (E), Ikzf1$^{+/-}$;Ikzf4$^{-/-}$ (F), Ikzf1$^{-/-}$;Ikzf4$^{+/-}$(G) or Ikzf1$^{-/-}$;Ikzf4$^{-/-}$ (H). Black arrows denote the fetal liver. RPL: Retinal progenitor layer. Scale bars: 10μm.

Figure S5, Javed et al.

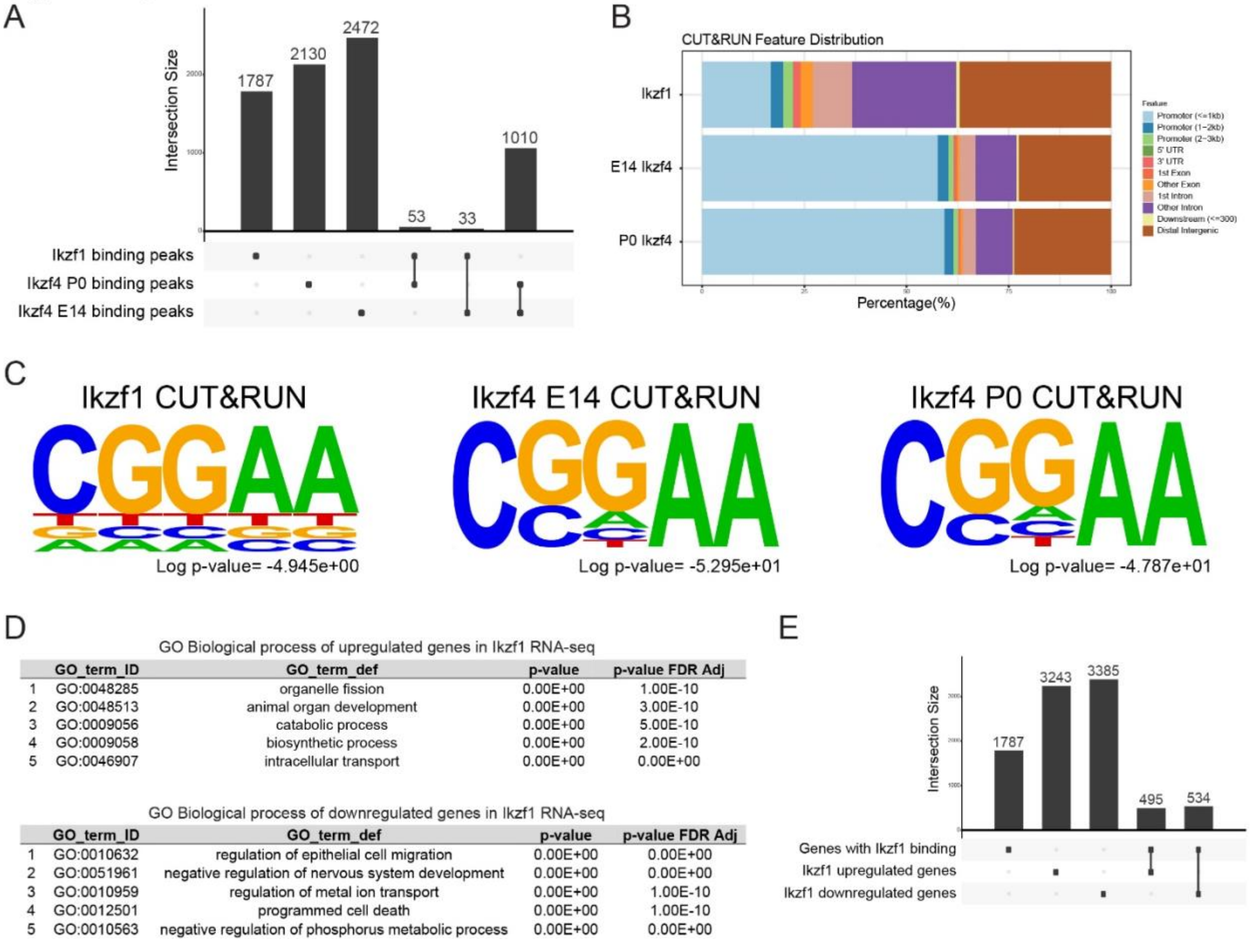


**Figure S5: Ikzf4 binds and upregulates Pou2f1/2 during retinal development and Müller glia specification genes during late retinogenesis.**

(A) Upset plot representing the overlap between Ikzf1 binding peaks and Ikzf4 E14/P0 binding peaks. (B) ChIP-seeker genomic annotation of the Ikzf1 and Ikzf4 E14/P0 CUT&RUN peaks across the entire genome. (C) HOMER analysis showing canonical 'GGAA' motif enriched in Ikzf1 and Ikzf4 E14/P0 CUT&RUN peaks. (D) GOnet analysis of all significantly up- and downregulated genes from DESeq2 analysis of Ikzf1 RNA-seq. (E) Upset plot showing gene overlap between Ikzf1 up/downregulated genes and genes bound by Ikzf1.

Figure S6, Javed et al.

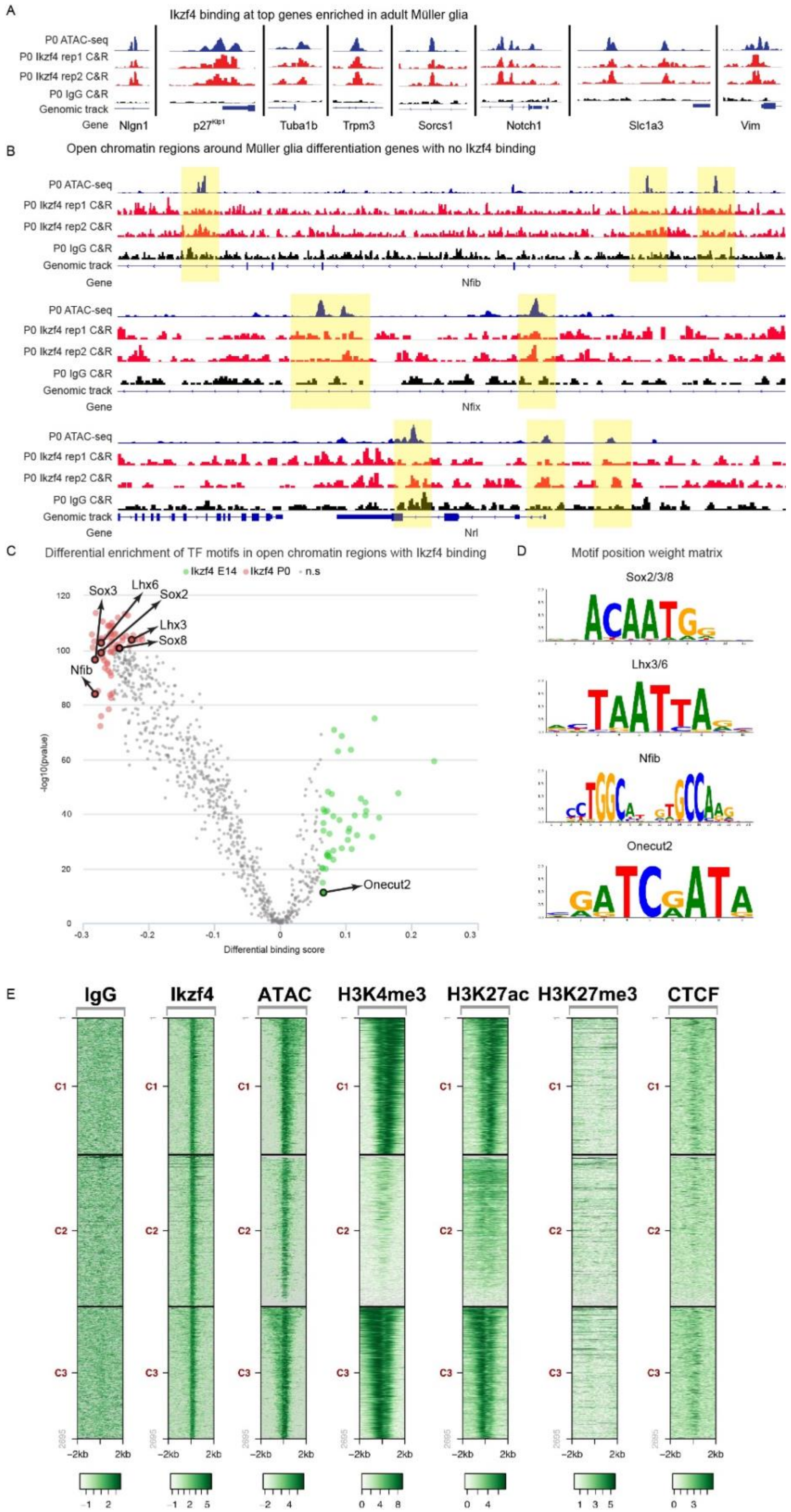

**Figure S6: Ikzf4 binds to cis regulatory regions important for Müller glia differentiation.** (A) Genomic tracks of ATAC-seq at P0 in blue (Aldiri et al. 2017), two replicates of Ikzf4 CUT&RUN at P0 in red, or IgG Control CUT&RUN at P0 in black at genomic location of Müller glia genes enriched in scRNA-seq dataset of P14 mouse retinas (Clark et al. 2019). (B) Genomic tracks of ATAC-seq at P0 in blue (Aldiri et al. 2017), two replicates of Ikzf4 CUT&RUN at P0 in red, or IgG Control CUT&RUN at P0 in black at genomic location of *Nfib*, *Nfix* and *Nrl*. Yellow highlighted areas indicate regions with positive ATAC-seq signal but negative Ikzf4 CUT&RUN signal. (C) Volcano plot of TOBIAS BINDetect TF footprinting analysis on open chromatin regions bound by Ikzf4 at E14 and P0. Arrows indicate motif with TF name. (D) Representative motif position weight matrix for TFs listed in (C). (E) Seqplot heatmaps of P0 IgG C&R, P0 Ikzf4 C&R, ATAC-seq, H3K4me3, H3K27ac, H3K27me3 and CTCF ChIP-seq from (Aldiri et al. 2017). TF: Transcription factor.

Figure S7, Javed et al.

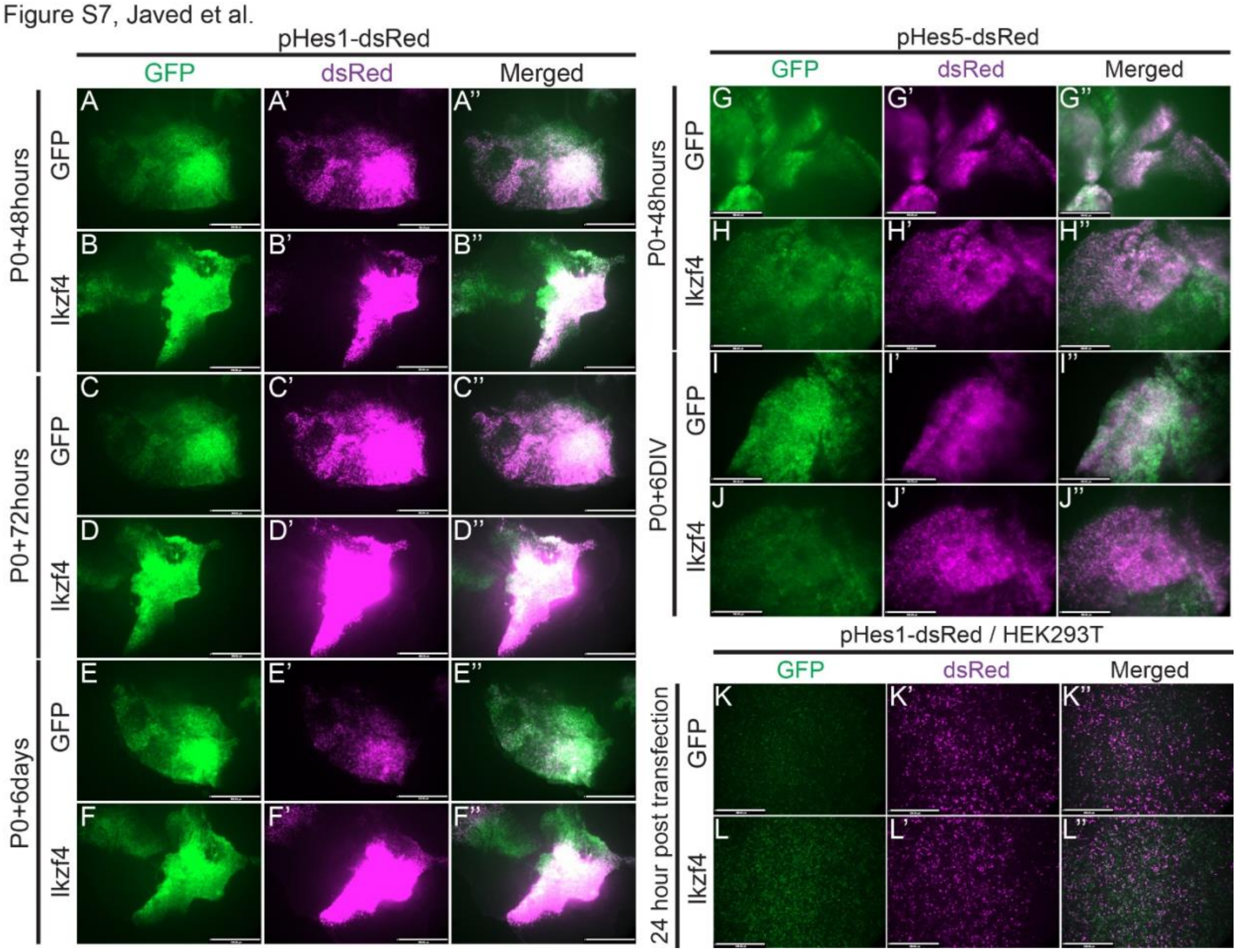


**Figure S7: Ikzf4 maintains Hes1 expression in post-mitotic cells during late retinogenesis.** (A-J'') Photomicrographs of retinal explants co-electroporated at P0 with Hes1-dsRed or Hes5-dsRed and either GFP (A-A'', C-C'', E-E'', G-G'', I-I'') or Ikzf4-IRES-GFP (B-B'', D-D'', F-F'', H-H'', J-J'') followed by imaging after either 48hours (A-B'', G-H''), 72hours (C-D''), or 6 days (E-F'', I-J''). (K-L'') Photomicrographs of HEK293T cells transfected with Hes1-dsRed and either GFP (K-K'') or Ikzf4-IRES-GFP (L-L'') imaged 24hours post-transfection. Scale bars: 360μm (A-L'').

**SUPPLEMENTARY FILES:**

**Supplementary file 1:** Excel spreadsheets containing the genes up/downregulated and bound by Ikzf1 compared to genes bound by Ikzf4 at E14 and P0 after GREAT analysis.

**Supplementary file 2:** Excel spreadsheet describing GOnet GO term annotation of the genes found in Supplementary file 1.

**Supplementary file 3:** Excel spreadsheet showing GREAT analysis of Ikzf4 E14 exclusive, P0 exclusive and Common binding target gene and peak location.

**Supplementary file 4:** Excel spreadsheet containing the complete list of differential enriched motifs between Ikzf4 E14 and Ikzf4 P0 open chromatin regions using TOBIAS BINDetect.

**Table S1: Sequences of primers.**

| mRNA primers | | | | |
|---|---|---|---|---|
| No. | | pF (5'>3') | pR (5'>3') | References |
| 1 | *β-actin* | TGATGGTGGGAATGG GTCAGAA | TCCATGTCGTCCCAGTTG GTAA | Batsché et al. 2005 |
| 2 | *Casz1v1* | CTTCGGGAACTGCAA GTACG | GTTGATGTGGTCCAAGCA GTG | Javed et al. 2020 |
| 3 | *Casz1v2* | TCGCAGAGTTACACTG GCTG | GGATCCCAACGGATCACT GG | Javed et al. 2020 |
| 4 | *Foxn4* | AATGATCAGAAGCTCG GGGC | CAGGACAGCGACTGAAGG TC | This paper |
| 5 | *Gapdh* | TGCAGTGGCAAAGTG GAGAT | ACTGTGCCGTTGAATTTGC C | Ouimette et al. 2010 |
| 6 | *Ldb1* | TTCTGCTTGGAGGATG GACC | ATGCTTCGGAAGTAGCGT GG | This paper |
| 7 | *Lhx2* | GGCAAGATCTCTGACC GCTA | TGCTGAAGCAGGTGAGTT CC | This paper |
| 8 | *Nfia* | GCATAGGGTGACAGC AACCA | ACCTAAACTGCCTTGGTCG G | This paper |
| 9 | *Nfib* | CTCATGAAGTCCCCGC ACTG | CTCCTGCACGTAGTATGC CAA | This paper |
| 10 | *Nfix* | AGCTTTCATCCCCGCT AAGG | TTGTAATGTCCTCGCGGCT C | This paper |
| 11 | *Nr2e3* | AAGCTCCTGTGTGACA TGTTCAA | AAGCTCCTGTGTGACATGT TCAA | Javed et al. 2020 |
| 12 | *Nrl* | CGAGCAGTGCACATCT CAGTTC | AACTGGAGGGCTGGGTTA CC | Javed et al. 2020 |
| 13 | *Pou2f2* | CACCACCAACAGCACA AACC | GGGGTTCAGGCCCGACAA G | Javed et al. 2020 |
| 14 | *Rnf12* | AATGGATCGCTTGGAT CGGG | GTACTTTCACCTGGGGTG CC | This paper |
| 15 | *Rxrg* | CCTCAATGCTCTTGGC TCTC | AGCTGCTGACACTGTTGA CC | This paper |
| 16 | *Sox8* | CGAGCAATGGAAGCA AGCAA | CTGAGCTCGGAGATGTCC AC | This paper |
| 17 | *Sox9* | CAAAACCGACGTGCAA GCTG | TCAGTTCACCGATGTCCAC G | This paper |

| Infusion HD sequences | | | | |
|---|---|---|---|---|
| 1 | Hes1-mut1-dsRed | CCGCGTGTCTGATATCC CCATTGGCTGAAA | TTTCAGCCAATGGGGATA TCAGACACGCGG | This paper |
| 2 | Hes1-mut2-dsRed | AAAGTTACTGTGATATCA AAGTTTGGGAA | TTCCCAAACTTTGATATC ACAGTAACTTT | This paper |

| | | | | |
|---|---|---|---|---|
| 3 | Hes1-mut3-dsRed | TGGGAAAGAAAGTTTGATATCTTTCACACGAGC | GCTCGTGTGAAAGATATCAAACTTTCTTTCCCA | This paper |
| 4 | Hes1-mut2+3-dsRed | GAAAGTTACTGTGATATCGAAAGTTTGATATCTTTCACACGAGCC | GGCTCGTGTGAAAGATATCAAACTTTCGATATCACAGTAACTTTC | This paper |

| shRNA sequences | | | | |
|---|---|---|---|---|
| 1 | shPou2f2 | gatccGCGCCAAATCTATTCCAGCTTTCAAGAGAAGCTGGAATAGATTTGGCGTTTTTTAAGCTTg | aattcAAGCTTAAAAAACGCCAAATCTATTCCAGCTTCTCTTGAAAGCTGGAATAGATTTGGCGCg | Javed et al. 2020 |

**Table S2 : Materials**

| REAGENT or RESOURCE | SOURCE | IDENTIFIER | APPLICATION |
|---|---|---|---|
| **Primary antibodies** | | | |
| Brn3a | Synaptic Systems | 411004 | IF 1:2000 |
| Brn3b | Santa Cruz | SC-6026 | IF 1:500 |
| Chx10 | Exalpha | X1180P | IF 1:500 |
| Crx/Otx2 | R&D systems | AF1979 | IF 1:500 |
| GFP | Abcam | ab13970 | IF 1:2000 |
| Hes1 | Cell Signaling Technology | 11988S | IF 1 :500 |
| Ikzf1 | Santa Cruz | M-20 (sc-9859) | IF 1 :100 |
| Ikzf4 | Millipore Sigma | ABE1331 | IF 1:100, C&R 2ug |
| Ki-67 | BD Biosciences | 550609 | IF 1:100 |
| Lhx2 | Fischer Scientific | PA5-78287 | IF 1 :200 |
| Lim1/2 | DSHB | 4F2 | IF 1:50 |
| M-opsin | Millipore Sigma | AB5405 | IF 1 :200 |
| Nfia/b | DSHB | 2C6 | IF 1 :500 |
| Nrl | R&D systems | AF2945 | IF 1:200 |
| Nr2e3 | Chemicon | Discontinued | IF 1 :200 |
| PNA-647 | Molecular Probes | L-32460 | IF 1:1000 |
| Pax6 | DSHB | AB_528427 | IF 1 :100 |
| Pax6 | Millipore Sigma | AB2237 | IF 1:100 |
| Rabbit control IgG Isotope | Fischer Scientific | 02-6102 | C&R 2ug |
| Rxrg | Abcam | AB15518 | IF 1 :200 |
| S-opsin | Santa Cruz | N-20 | IF 1:1000 |
| Sox2 | Abcam | AB97959 | IF 1 :200 |
| **Secondary antibodies** | | | |
| AF-488 Donkey anti-Chicken | Jackson ImmunoResearch | AB_2340375 | IF 1:1000 |
| AF-488 Donkey anti-Mouse | Jackson ImmunoResearch | AB_2340846 | IF 1:1000 |
| AF-488 Donkey anti-Guinea Pig | Jackson ImmunoResearch | AB_2340472 | IF 1:1000 |
| AF-555 Donkey anti-Rabbit | Jackson ImmunoResearch | AB_2313584 | IF 1:1000 |
| AF-647 Donkey anti-Goat | Jackson ImmunoResearch | AB_2340436 | IF 1:1000 |
| AF-647 Donkey anti-Mouse | Jackson ImmunoResearch | AB_2340863 | IF 1:1000 |
| AF-647 Donkey anti-Sheep | Jackson ImmunoResearch | AB_2340751 | IF 1:1000 |
| **Bacterial and Virus Strains** | | | |
| Subcloning Efficiency™ DH5α™ Competent Cells | Fischer Scientific | 18265017 | |
| **Chemicals, Peptides, and Recombinant Proteins** | | | |
| Papain | Worthington | LS003124 | |

| | | |
|---|---|---|
| **Critical Commercial Assays** | | |
| RNeasy Microkit | Qiagen | 74004 |
| Superscript VILO Master Mix | Fisher Scientific | 11755050 |
| SYBR Green Master mix | Fisher Scientific | A25742 |
| Dynabeads Protein G | Fisher Scientific | 10003D |
| In-Fusion HD Cloning plus | Takara | 638920 |
| Click-iT™ EdU Alexa Fluor™ 647 | Fisher Scientific | C10340 |
| **Experimental Models: Cell Lines** | | |
| Phoenix-AMPHO | ATCC | CRL-3213 |
| **Experimental Models: Organisms/Strains** | | |
| CD1 | Charles Rivers | Cat#022 |
| Ikzf1$^{-/-}$ | Wang et al. 1996 | N/A |
| Ikzf4$^{-/-}$ | International Mouse Phenotyping Consortium and RIKEN Bioresource | RBRC06808 |
| **Oligonucleotides** | | |
| For qPCR primers, see Table S1 | This paper | N/A |
| For Infusion HD sequences, see Table S1 | This paper | N/A |
| **Recombinant DNA** | | |
| pCIG2-Ikzf1-IRES-GFP | Mattar et al. 2015 | N/A |
| pCIG2-Ikzf4-IRES-GFP | This paper | N/A |
| pCLE-venus | Addgene Gaiano et al. 2000 | 17703 |
| pCLE-Ikzf4-IRES-venus | This paper | N/A |
| pCIG2-IRES-GFP | Hand et al. 2005 | N/A |
| pHes1-dsRed | Addgene Matsuda and Cepko. 2007 | 13767 |
| pSIREN-RetroQ-ZsGreen | Clontech | 632455 |
| **Software** | | |
| Prism 8 | Graphpad | https://www.graphpad.com/scientific-software/prism/ |
| Volocity software 6 | Improvision | http://www.perkinelmer.com/lab-solutions/resources/docs/BRO_VolocityBrochure_PerkinElmer.pdf; |

| | | |
|---|---|---|
| ZEN software | Zeiss Microscope | https://www.zeiss.com/microscopy/int/products/microscope-software/zen.html |
| Adobe Illustrator CC 2020 | Adobe | http://www.adobe.com/products/illustrator.html |
| Adobe Photoshop CC 2020 | Adobe | https://www.adobe.com/products/photoshop.html |
| Adobe Acrobat Pro DC 2020 | Adobe | https://acrobat.adobe.com/us/en/acrobat.html |
| Quant-Studio Real Time PCR software | Fisher Scientific | https://www.thermofisher.com/ca/en/home/life-science/pcr/real-time-pcr/real-time-pcr-instruments/quantstudio-3-5-real-time-pcr-system.html |
| Office 365 | Microsoft | https://www.office.com/ |
| Cellranger (4.0) | 10x Genomics | https://support.10xgenomics.com/single-cell-gene-expression/software/overview/welcome |
| RStudio (1.3.1056) | Rstudio | https://rstudio.com/ |
| R (4.0.2) | The R Project for Statistical Computing | https://www.r-project.org/ |
| TOBIAS (0.13.0) | Bentsen et al. 2020 | https://github.com/loosolab/TOBIAS |
| Seurat (3.2.1) | Butler et al. 2018 | https://satijalab.org/seurat/ |
| Spyder (4.1.4) | Spyder IDE | https://www.spyder-ide.org/ |
| Python (3.8.3) | Python programming language | https://www.python.org/ |
| Scanpy (2.1.0) | Wolf et al. 2018 | https://scanpy.readthedocs.io/en/stable/index.html |
| Velocyto.py (0.17) | La Manno et al. 2018 | http://velocyto.org/velocyto.py/index.html |
| Galaxy platform | Afgan et al. 2016 | https://usegalaxy.org/ |
| Bowtie2 (2.3.4.3) | Langmead and Salzberg et al. 2012 | http://bowtie-bio.sourceforge.net/bowtie2/index.shtml |
| MACS2 (2.1.1.20160309.6) | Feng et al. 2012 | https://github.com/taoliu/MACS/releases |
| Bedtools (2.28.0) | Quinlan and Hall. 2010 | https://bedtools.readthedocs.io/en/latest/content/bedtools-suite.html |

| | | |
|---|---|---|
| Deeptools2 (3.3.2.0.0) | Ramirez et al. 2016 | https://deeptools.readthedocs.io/en/develop/ |
| Integrative Genomics Viewer (IGV) (2.8.13) | Broad Institute | http://software.broadinstitute.org/software/igv/ |
| Biorender | Biorender | https://biorender.com/ |
| **Other** | | |
| DMEM+Glutamax | Thermofisher | 10569010 |
| Penicillin/Streptomycin | Thermofisher | 15140148 |
| Fetal Bovine Serum | Wisent Bioproducts | 080-150 |
